# A QbD strategy to develop curcumin and siRNA co-loaded lipoplexes to combat osteoarthritis-related inflammation and oxidative stress

**DOI:** 10.1101/2024.11.25.625050

**Authors:** Saketh Reddy Ranamalla, Alina Porfire, Emilia Licarete, Lucia Tefas, Rohith Pavan Parvathaneni, Oommen. P. Varghese, Alina Sesarman, Monica Focsan, Lucian Barbu Tudoran, Ioan Tomuta, Manuela Banciu

## Abstract

Chronic knee and lower back pain due to osteoarthritis (OA) has a global prevalence and impacts human well-being by impairing mobility. Oxidative stress and inflammation are key factors in OA pathogenesis and progression. Non-viral gene delivery through liposomes is a promising approach for repairing damaged cartilage tissues. Our study focuses on developing co-loaded lipoplexes for efficient co-delivery of curcumin and therapeutic siRNA. Curcumin downregulates many inflammatory cytokines, scavenges free radicals, and upregulates collagen and aggrecan, therefore reducing pain and helping with regeneration. Quality by Design (QbD) principles guided the development of curcumin-loaded cationic liposomes (CL), which were further used as vectors for therapeutic siRNA in the design of the co-loaded lipoplexes. QbD steps involved risk assessment, Design of Experiments (DoE), and selection of the optimal vector, i.e., optimum curcumin-loaded cationic liposomes (Opt-CL), which would ensure the best transfection efficiency for therapeutic siRNA. The efficiency of Opt-CL was evaluated in both the primary chondrocytes and cell lines, which were induced by oxidative stress and inflammatory conditions. The Opt-CL successfully reduced the oxidative stress levels in both. The Opt-CL were complexed with the IL-6 and IL-8 siRNA to form co-loaded lipoplexes, which efficiently reduced the inflammation in the chondrocytes. These co-loaded lipoplexes effectively transfected chondrocytes with no toxicity and are promising for further testing in OA models. The study has yielded an optimal non-viral vector that could serve as a platform for the incorporation of other lipophilic drugs and negatively charged oligonucleotides for various ailments.

**Highlights:**

- Utilization of QbD to screen and optimize critical factors for the development of CL
- Development of proof of concept using the curcumin and luciferase siRNA as the small molecule and oligonucleotide candidates for efficient cell viability and transfection
- Evaluation of cell internalization and gene knockdown in a luciferase-expressing chondrocyte cell line to prove the model efficacy
- In chondrocytes, the optimized formulation demonstrated the reduction of inflammation and oxidative stress.

## 1. Introduction

Osteoarthritis (OA) remains a prevalent and debilitating disease characterized by joint degeneration, pain, and loss of mobility. Despite considerable research, the treatments available have mainly focused on symptom management rather than addressing the underlying pathogenesis of the disease [1]. The pathogenesis of OA involves complex biological processes, including mechanical stress, inflammation, and oxidative stress, which interplay to drive disease progression. Inflammation is a primary characteristic of OA, with pro-inflammatory cytokines such as interleukin-1β (IL-1β) and tumor necrosis factor-alpha (TNF-α) playing pivotal roles. These cytokines are produced by chondrocytes, synovial cells, and infiltrating macrophages, leading to the upregulation of matrix metalloproteinases (MMPs) and aggrecanases (ADAMTS 4 and 5), which degrade cartilage extracellular matrix components like collagen and aggrecan [2]. Studies have shown that in IL-1 β-induced chondrocytes, the essential mediators involved in OA pathogenesis are MMP −1,3 and 13, Interleukins (IL) like IL-1β, IL-6, IL-8, IL-17. MMPs jointly act to destroy essential structural components of the cartilage, interstitial collagen, proteoglycans, fibronectin, type IX collagen, and type II collagen [3]. At the same time, IL-6 is a multifunctional cytokine that stimulates the production of MMPs, such as MMP-1, MMP-3, and MMP-13, and other inflammatory mediators, exacerbating cartilage breakdown. IL-8 and IL-17 amplify the inflammatory response by enhancing the expression of IL-1β and TNF-α [4]. Furthermore, oxidative stress, via reactive oxygen species (ROS), activates signaling pathways such as Nuclear Factor kappa-light-chain-enhancer of activated B cells (NF-κB) and Mitogen-Activated Protein Kinase (MAPK), worsening cartilage damage [5]. This dysregulated inflammatory tissue environment in OA represents a therapeutic target, as modulating the activity of pro-inflammatory cytokines, MMPs, and oxidative stress pathways such as NF-κB and MAPK could potentially halt or reverse cartilage degradation.

Given this complex pathogenesis, this study aimed to develop a novel therapeutic strategy that combines curcumin with gene-silencing small interfering RNA (siRNA) technology to address multiple aspects of OA progression at the molecular level. Curcumin, derived from turmeric, has been identified as a potent anti-inflammatory and antioxidant agent with the potential to alleviate OA symptoms and even slow disease progression [6,7]. Its anti-inflammatory effects stem from inhibiting the NF-κB pathway, which regulates pro-inflammatory cytokines like IL-1β, IL-6, IL-8, and IL-17. Curcumin blocks NF-κB activation by preventing Inhibitor of kappa B (IκB) degradation, reducing the expression of cytokines and enzymes like Cyclooxygenase-2 (COX-2) and Inducible Nitric Oxide Synthase (iNOS) [8]. It also inhibits other inflammatory pathways, including MAPK and Janus Kinase/Signal Transducer and Activator of Transcription (JAK/STAT), thereby limiting the inflammatory response [9]. Curcumin’s antioxidant properties mitigate oxidative stress by scavenging ROS [10] and enhancing endogenous antioxidant enzymes such as superoxide dismutase (SOD), catalase, and glutathione peroxidase [7]. It also inhibits ROS-generating enzymes like NADPH oxidase [11], reducing oxidative damage and inflammatory signaling. Despite its therapeutic potential, curcumin has poor oral bioavailability and consequently has limited clinical application. To overcome the poor bio-availability, local intra-articular delivery based on liposomes [12] shows promise for improving curcumin’s effectiveness in OA treatment [13].

Nevertheless, the complexity of OA’s pathogenesis, involving cartilage wear, inflammation, and bone remodeling, necessitates innovative therapeutic strategies targeting multiple aspects of the disease process [2]. Thus, developing a therapy that exploits combinations of small-molecule drugs, such as curcumin, with gene-silencing drugs, such as small interfering RNA (siRNA), offers a novel approach to OA treatment [14]. Since siRNA-based drugs can silence the expression of any disease-causing genes at the post-transcriptional level, this strategy offers an excellent opportunity to potentially halt the destruction of osteoarticular tissue. By silencing mRNAs for specific genes encoding cytokines involved in the OA pathogenic process, siRNA therapy has the potential to modulate disease progression at the molecular level [15]. However, efficient intracellular siRNA delivery presents a significant challenge, namely the risk of compromising the structural integrity of the molecule in the biological environment. One of the ways to overcome this challenge is to use cationic liposomes that facilitate efficient encapsulation and protection against nucleases, thereby enhancing their delivery and efficacy in target cells [16].

Cationic liposomes are promising for delivering therapeutic agents and nucleic acids directly to disease sites. Their ability to encapsulate both hydrophobic and hydrophilic drugs makes them suitable for delivering anti-inflammatory agents, cartilage-repair factors, and gene-modulating material [17–20]. These nanocarriers offer advantages such as enhanced stability, controlled release, and biocompatibility [16]. Given the complexity of their development, the QbD approach represents an efficient way to obtain nanocarriers with optimal quality characteristics.

Implementing QbD in developing cationic liposome formulations ensures product quality through careful manufacturing process design rather than relying solely on post-production testing. This approach has proven effective in optimizing nanoparticle-based drug delivery systems by systematically exploring formulation variables to achieve optimal particle size, charge, encapsulation efficiency, and release profiles [21]. Regulatory agencies like the European Medicines Agency (EMA) and Food and Drug Administration (FDA) promote adherence to ICH guideline Q8(R2), which outlines QbD principles. A QbD study typically involves A) defining the Quality Target Product Profile (QTPP), B) identifying critical quality attributes (CQAs), material attributes (CMAs), and process parameters (CPPs), C) conducting risk assessment and design of experiments (DoE) to link CMAs and CPPs to CQAs, and D) establishing a design space, through formulation and process optimization [22].

This study presents a promising strategy for addressing OA’s therapeutic challenges using lipoplexes co-loaded with curcumin and siRNA. By applying QbD principles, the curcumin-loaded liposomal formulations were prepared and were evaluated for their physicochemical properties. These formulations were then loaded with the luciferase siRNA to produce the co-loaded lipoplexes. *In vitro*, all the formulations were assessed for cell viability and transfection efficacy using the C28/I2 chondrocyte cell line, modified to express firefly luciferase. Successful knockdown of luciferase expression with the designed siRNA validated efficient oligonucleotide delivery and mRNA suppression. These responses’ data helped us establish a design space, guiding the preparation of optimized formulations. Furthermore, the optimized formulations demonstrated curcumin’s antioxidant and anti-inflammatory effects and siRNA-mediated reductions in IL-6 and IL-8 cytokine levels, confirming the therapeutic potential of this approach. The combined anti-inflammatory, antioxidant, and chondroprotective effects of curcumin, combined with siRNA’s gene-silencing capabilities, offer a targeted, multifunctional OA therapy.

## 2. Materials and methods

### 2.1. Materials

For oligonucleotide synthesis (procedure in the supplementary file, MALDI TOF analysis in Figure S.1), acetonitrile, ammonium hydroxide solution, methylamine solution (40% in water), triethylamine trihydrofluoride, isopropoxy trimethyl silane, diethyl ether, dimethyl sulfoxide, and urea were purchased from Sigma-Aldrich (Saint Louis, USA). Universal UnyLinker Support 1000 Å, 2’-O-TBDMS protected nucleoside phosphoramidites, DMT removal reagent (3% trichloroacetic acid/DCM), activation reagent (0.3M benzyl thiotetrazole/acetonitrile), CAP A (acetic anhydride/pyridine/THF), CAP B (16% N-methylimidazole in THF), oxidation solution (0.02M iodine/pyridine/water/THF) were purchased from ChemGenes (Wilmington, USA). For liposomes, curcumin was obtained from Sigma-Aldrich (Saint Louis, USA), 1,2-Dioleoyloxy-3-trimethylammonium-propane chloride (DOTAP), 1,2-Dipalmitoyl-sn-glycero-3-phosphocholine (DPPC), 1,2-Dioleoyl-sn-glycero-3-phosphocholine (DOPC), 1,2-Dioleoyl-sn-glycero-3-phosphoethanolamine (DOPE) and N-(carbonyl-methoxy polyethyleneglycol-2000)-1,2-distearoyl-sn-glycero-3-phosphoethanolamine (Na-salt; MPEG-2000-DSPE) were obtained from Lipoid GmbH (Ludwigshafen, Germany); cholesterol (CHO) was obtained from Sigma-Aldrich (Saint Louis, USA); Spectra/Por 3 Dialysis Tubing, 3.5 kDa MWCO for release studies was obtained from Repligen (Massachusetts, USA). For Lipoplexes, IL-6 siRNA (Cat.No: 4390824 - s7313), IL-8 siRNA (Cat.No: 4390824 - s7328), Scramble (negative control) siRNA (Cat.No: 4390843) were obtained from ThermoFisher Scientific (Eugene, USA). The nucleic acid stain SYBR Gold (ThermoFisher Scientific, USA), Agarose, and gel loading buffer (Sigma-Aldrich, USA) were obtained for complexation capacity. The alamarBlue™ Cell Viability Reagent was used for cell viability and purchased from ThermoFisher Scientific (Eugene, USA). Luciferase assay was performed using the ONE-Glo Luciferase assay system from Promega (Madison, USA) to quantify the cell transfection. For nucleic acid quantification, Quant-iT™ PicoGreen™ dsDNA Assay Kits and dsDNA Reagents were used, which were obtained from ThermoFisher Scientific (Eugene, USA). The qRT-PCR kits were purchased from BioRad (California, USA), and the ELISA kits were purchased from Peprotech ThermoFisher Scientific (Massachusetts, USA). Fluorescent Universal Negative Control #1, Cyanine 5 (# SIC005) was obtained from Sigma (Saint Louis, USA) for confocal microscopy.

### 2.2. Preparation and evaluation of curcumin-loaded cationic liposomes (CL)

The CLs were developed following the QbD approach and were evaluated through *in vitro* cell studies. This approach has helped us select the optimum curcumin-loaded cationic liposomes (Opt-CL), which were further evaluated for stability, release profile, and antioxidant effect.

#### 2.2.1. QbD-based development of CL

Developing cationic liposomes involves many variables and requires a systematic and quality-driven approach. So, in this study, we employed QbD principles to ensure the robustness and reproducibility of the formulation process.

##### 2.2.1.1. QTPP and Risk Assessment

Establishing a QTPP was a crucial foundation for identifying CQAs and defining the desired characteristics of the final product. The QTPP provides clear objectives for the formulation development process, including the desired therapeutic efficacy, stability, and safety. In addition to establishing the QTPP, we utilized the Ishikawa diagram, also known as the fishbone diagram, which helped to visually map out potential causes contributing to identified failure modes, providing a comprehensive understanding of risks in the formulation process before proceeding to the Risk assessment. A Failure Mode and Effects Analysis (FMEA) was pivotal in identifying potential risks associated with the formulation process and guiding risk mitigation strategies. It is a systematic method for identifying potential failure modes within a process, assessing each failure mode’s severity, occurrence, and detectability (each on a scale of 1 to 5), and prioritizing them based on their Risk Priority Number (RPN) given in equation 1 [23].

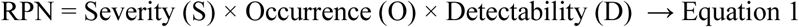

##### 2.2.1.2. DoE and the design space

A screening study was used to evaluate all the potential risk variables and give us a comprehensive understanding of the formulation space. The experiments were designed, and the data were statistically analyzed using MODDE 13 Pro software (Umetrics, Sartorius, Malmö, Sweden). Four variables were established as independent factors to be investigated, namely the helper lipid type (X1), the concentration of cationic lipid (X2), the concentration of cholesterol (X3), and the concentration of pegylated phospholipid (X4). Each independent factor further had three more levels. Based on these variables and their levels, an L18 fractional factorial design with 18 runs and 3 center points for a total of 21 runs was considered suitable. The investigated responses were the CQAs of the cationic liposomes like the particle size (Y1), PDI (Y2), zeta potential (Y3), curcumin entrapment efficiency (Y4), siRNA complexation capacity (Y5), fluorescence level (Y6), cell viability (Y7), and cell transfection (Y8). The data were fitted by means of partial least squares using the statistical module of MODDE 13 Pro software. The experiments were performed in a random order to reduce the experimental variability. This design allowed us to screen all the potential risk variables, gave us a comprehensive understanding of the formulation space, and helped identify the design space for the preparation of the Opt-CL.

#### 2.2.2. Preparation of CL

CL was prepared by using a modified thin film hydration technique. All lipids and curcumin were weighed accurately according to the concentrations provided in the DoE and dissolved in ethanol. The resulting solution was transferred into a round-bottom flask (RBF), and the solvent was evaporated in a rotary evaporator at 45°C temperature and 100 rpm. Once the lipid film was formed inside the RBF, it was hydrated by adding PBS and sonicated until it was wholly dispersed in water. The dispersion was subsequently extruded through 800 nm, 600 nm, 400 nm, and 200 nm polycarbonate membranes, five times each, by maintaining the temperature of the LiposoFast LF-50 extruder from Avestin Europe GmbH (Mannheim, Germany) at 45°C. Finally, the extruded liposomes were physicochemically characterized and evaluated *in vitro*.

#### 2.2.3. Characterization of CL

The extruded curcumin-loaded CL were characterized for particle size (PS), polydispersity index (PDI), zeta potential (ZP), encapsulation efficiency (EE), complexation capacity (CC), and *in vitro* tests.

##### 2.2.3.1. Particle size, PDI, and Zeta potential

The CL obtained were diluted 100 times using distilled water and were evaluated for PS, PDI, and ZP using the Malvern Zetasizer Nano ZS (Worcestershire, UK). The dynamic light scattering principle determined the PS and PDI at an angle of 90° at 25°C. The ZP was determined by the electrophoretic mobility of the liposomes at 25°C.

##### 2.2.3.2. Encapsulation efficiency

The EE of curcumin in CL was evaluated by HPLC using the method reported by Tefas et al [24]. Firstly, the liposomes were centrifuged at 2300 g for 30 minutes to sediment the unentrapped curcumin. The liposomes in the supernatant with entrapped curcumin were collected. To quantify the entrapped curcumin, one part of the liposomes was mixed with nine parts of ethanol, which solubilized the lipids and the curcumin. The chromatographic separation was performed on an Agilent 1100 Series HPLC system (Agilent Technologies, CA, USA) equipped with a Zorbax SB C18 column. The injection volume was 5 uL, and the mobile phase consisted of acetonitrile and 0.2% formic acid (35:65) at a flow rate of 1 mL/min and 30 °C. The UV detection wavelength was set at 420 nm. All measurements were performed in triplicates. The entrapment efficiency was calculated by equation 2.

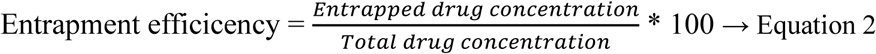

##### 2.2.3.3. Complexation capacity

The CL obtained through DoE was checked for siRNA complexation capacity at different Nitrogen to Phosphate (N/P) ratios. Briefly, the previously prepared positively charged liposomes were mixed with negatively charged luciferase siRNA (preparation procedure in Supplementary file) for 30 minutes in an Eppendorf tube using a thermal mixer from Thermo Scientific™ (Massachusetts, USA) in N/P ratios of 2.5:1, 5:1, 10:1 and 20:1. After mixing, the resulting lipoplexes were put aside for another 30 minutes and subsequently evaluated as described further on.

The uncomplexed free nucleic acid was separated from the lipoplexes using agarose gel electrophoresis. A reference sample of free nucleic acid was run on the same gel to serve as a control. The nucleic acid bands were then visualized using the nucleic acid stain SYBR Gold, which binds non-selectively to nucleic acids and was imaged by the ChemiDoc Imaging System from Bio-Rad (California, USA). The bands were subsequently quantified using ImageJ software. The complexation capacity is calculated by comparing the uncomplexed nucleic acid from the lipoplexes to the reference of the free nucleic acids, using the following equation 3,

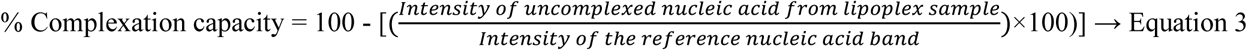

#### 2.2.4. *In vitro* evaluation of CL

The luciferase-expressing C28/I2 cells were cultivated to evaluate the performance of the developed liposomes, i.e., their ability to be internalized by the cells and to transfer the complexed nucleic acids into the cells. All the CL formulations were evaluated were evaluated for cell viability by resazurin assay, liposome cell internalization by fluorescence levels, and cell transfection by luciferase gene knockdown.

##### 2.2.4.1. Cell culture

The human C28/I2 chondrocyte cell line was cultured in T-75 flasks with DMEM GlutaMAX^TM^ medium from Gibco, ThermoFisher (Massachusetts, USA), at 37 °C, 5% CO_2_ and 80% humidity. The cells were passaged when they reached about 80% confluency. For the cell transfection and cell viability studies, the cells were seeded at a density of 5,000 cells per well in 96-well culture plates.

##### 2.2.4.2. Cell viability

Based on preliminary results, 50 μM curcumin-equivalent concentration of liposomes was tested on the cells. The influence of CL on the viability of C28/I2 cells was checked using the AlamarBlue assay, in which resazurin, a cell-permeable compound, is reduced to resorufin, a red-colored and highly fluorescent compound. The absorbance was checked using the microplate reader FLUOstar Omega from BMG Labtech (Ortenberg, Germany) [25]. at 570 and 600 nm. Cells untreated with liposomes were taken as control. The cell viability was calculated by using the following equation,

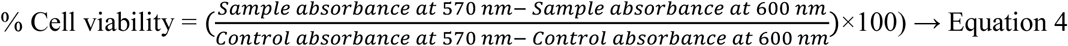

##### 2.2.4.3. Cell internalization

Cells were cultured in black 96-well plates compatible with fluorescence microscopy. The CL was added to the cell culture medium in curcumin-equivalent concentration of 50 μM for all the DoE formulations. Cells without the liposome treatment were considered as control. The cells were incubated for 24 hours and then washed thrice with sterile PBS and examined under the Zeiss Axio Vert.A1 microscope (Oberkochen, Germany) with a fluorescence detector, and images were captured. Additionally, the fluorescence was measured by using a plate reader FluoSTAR, BMG Labtech (Ortenberg, Germany) to assess the extent of liposomal uptake. The fluorescence intensity was calculated by using the following equation,

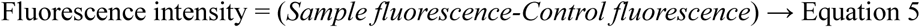

##### 2.2.4.4. Cell transfection efficiency studies

Based on the previous complexation data, a 20:1 N/P ratio for the liposomes was identified as optimal to ensure the complete complexation of 50 nM luciferase siRNA. Untreated cells were taken as control. The cells previously plated in 96-well plates were washed with PBS and transfected by adding the luciferase siRNA co-loaded lipoplexes mixed with OptiMEM medium. The cells were then incubated for 48 hours and lysed using RIPA lysis buffer. A part of the cell lysate was transferred onto the white plate to measure the chemiluminescence using the Luciferase Assay System. The relative light units (RLU) per well were determined using chemiluminescence according to the manufacturer’s instructions. The second part of the cell lysate was used for nucleic acid (NA) quantification using the Quant-iT Picogreen assay according to the manufacturer’s instructions. The cell transfection measured by the gene knockdown correlates with the reduction of the chemiluminescence compared to that of the non-transfected cells. The Lipofectamine RNAimax was used as a reference to evaluate the transfection efficiency of the liposomal formulations. The amount of chemiluminescence from the wells was normalized by nucleic acid content. The cell transfection (gene knockdown) was determined by equation 6.

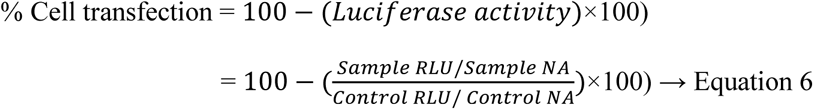

#### 2.2.5. Lyophilization, stability, and release profile of the Opt-CL

The Opt-CL formulation identified based on statistical evaluation of the results was evaluated for the physicochemical properties (PS, PDI, ZP, EE), in vitro behavior on cell line (CV and CT) and also further studied for stability, release profile, and cellular uptake at the subcellular level.

The Opt-CL was monitored for stability at 4°C for over six months. The physicochemical properties of the liposomes, i.e., the PS, PDI, ZP, and encapsulated curcumin retention (ECR), were evaluated with periodic sampling. The Opt-CL were lyophilized using 100 mM trehalose and mannitol mixture (1:5 molar ratio) as cryoprotectants. The liposomes were characterized for PS, PDI, and ZP before and after lyophilization (resuspended formulation). Transmission Electron Microscopy (TEM) was performed to confirm the morphological integrity of the Opt-CL and the blank liposomes (same lipid composition as Opt-CL without curcumin) before and after lyophilization. Liposomes were diluted with sterile water, and then the salts of PBS were removed using the Amicon Ultra-2 centrifugal filter units from Merck (New Jersey, USA). The sample was added on a formvar-coated carbon grid, followed by negative staining with uranyl acetate. After drying, the grids were imaged with a Jeol JEM 1010 instrument and a Mega View III CCD camera for image capturing (Tokyo, Japan).

The release profile of curcumin from the liposomes was tested by utilizing dialysis membranes with 3.5 kDa MWCO in a Phoenix™ diffusion cell apparatus from Teledyne Hanson Research (California, USA). These Franz diffusion cells, described by Siefan et al., were the closest replicable instruments for knee intra-articular space with a 10 mL receptor chamber and liposomes-filled donor chamber divided by the 3.5 kDa membrane [25]. The sink conditions were ensured, and the release profile for curcumin (Figure 4) was performed in PBS at pH 5, mimicking the late endosomal pH and PBS at pH 6.8, representing the inflamed joint environment [26,27]. Tween 80 was added as a solubilizer to the aqueous media to aid the release of lipophilic curcumin and also gives insight into mimicking the environment in the presence of factors that break down the liposomes [28–30]. The release media were maintained at 37°C and stirred at 500 rpm throughout the study. An autosampler regularly collected 0.25 mL samples from the receptor compartment at 0.5, 1, 2, 4, 8, 16, 20, 24, 36, 48, 60 and 72 hours. The samples were analyzed for the curcumin content using the HPLC method mentioned in the previous subsection.

For confocal microscopy, the liposomes without curcumin were prepared and complexed with Cyanine 5 (Cy5)-labeled siRNA (Sigma Aldrich, Missouri, USA). Cells were seeded onto µ-Dish 35 mm Petri dishes (Ibidi, Gräfelfing, Germany) and transfected with the Cy5-siRNA loaded lipoplexes for 48 hours. Free siRNA was used as a control in a separate dish. Post-transfection, cells were washed with PBS, fixed with 4% paraformaldehyde for 30 minutes, and permeabilized with 1% Triton X-100 for another 30 minutes. Cells were then stained with 1 μg/mL DAPI solution to visualize nuclei. The samples were imaged using a super-resolution Re-Scan Confocal Microscopy system (RCM-VIS unit) from Confocal.nl (Amsterdam, The Netherlands) equipped with two diode lasers (TOPTICA Photonics AG, Germany), 405 nm (RCM 405) and 561 nm (RCM 561), mounted on an ECLIPSE Ti2-E inverted microscope (Nikon). The RCM images were captured using a PCO EDGE 4.2 CCD camera and analyzed with NIS Elements Software to assess intracellular localization and distribution of the labeled liposomes.

#### 2.2.6. Assessment of the anti-inflammatory effect

##### 2.2.6.1. Cell culture and seeding

The luciferase-expressing C28/I2 cells and the human articular chondrocytes were cultivated to evaluate the anti-inflammatory and antioxidant potential of lipoplexes *in vitro*.

Human articular chondrocytes were received from patients undergoing knee surgeries at UMC, Utrecht. The isolation procedure of articular cartilage from 3 patients (all female, age: 61.5 ± 2) with OA undergoing total knee arthroplasty was previously reported by Katrin et al. The batch numbers C28-CA-22-004, C28-CA-22-009, and C28-CA-23-004 represent the patient-primary cells used. Cells were expanded until the first passage and frozen. Cells from passages two to four were used for the experiments [26]. The human OA chondrocytes (Passages 2 - 4) were seeded in well plates containing the chondrogenic medium (DMEM(1X) + GlutaMAX^TM^ (+ 4.5g/L D-glucose, + Pyruvate) supplemented with 0.2 nM ascorbic-2-phosphate, 1x ITS-X (10 mg/L insulin, 5.5 mg/L transferrin, 6.7 μg/L sodium selenite, two mg/L ethanolamine) (Thermo Fisher Scientific), 4.0 g/L human serum albumin (Sanquin), and 100 U/ml penicillin/streptomycin) with two mM HEPES (Gibco), for 24 hours.

##### 2.2.6.2. Inflammatory cytokine profiling

The primary aim of cytokine profiling was to identify the most commonly overexpressed cytokines in order to select and target them with their specific siRNA. qRT-PCR was used to determine the cytokine profile in IL-1β induced primary chondrocytes and the C28/I2 chondrocyte cell line. Therefore, the cells were seeded in a 60 mm sterile petri dish at a seeding density of about 80,000 cells per well. At around 50% confluency, the cells were treated with 10 ng/mL of IL-1β for 24 hours to create inflammatory conditions [27]. The total RNA from the lysed chondrocytes was isolated and purified using an RNeasy Mini RNA isolation kit from Qiagen (Hilden, Germany). The RNA obtained was quantified using the NanoDrop from ThermoFisher (Massachusetts, USA). cDNA was synthesized from a reaction mixture containing the RNA template and the supermix with oligonucleotides (iScript reverse transcription Supermix for qRT-PCR, BioRad, USA) following the manufacturer’s instructions. An appropriate quantity of cDNA was mixed with forward and reverse primers and a mixture of Sso7d-fusion polymerase and EvaGreen dye (Ssofast EvaGreen Supermix, BioRad, USA). The reaction was carried out on Bio-Rad CFX96 real-time PCR systems under the following parameters: enzyme activation at 95 °C for 30 seconds, 40 cycles (of denaturation, annealing/extension) at 95 °C for 5 seconds, 60°C for 5 seconds, and then melt curve from 65 – 95 °C (in 0.5°C increments) for 5 seconds per step. All the melting curves were generated to check for primer specificity. The cytokines’ primers checked are given in Table 1, where human β-actin was taken as a reference for gene expression. Comparative Ct method (ΔΔCt) was used to calculate gene expression. Gene expression was reported as fold change (2-ΔΔCt) relative to mRNA expression in inflammation-induced chondrocytes.

**Table 1.**
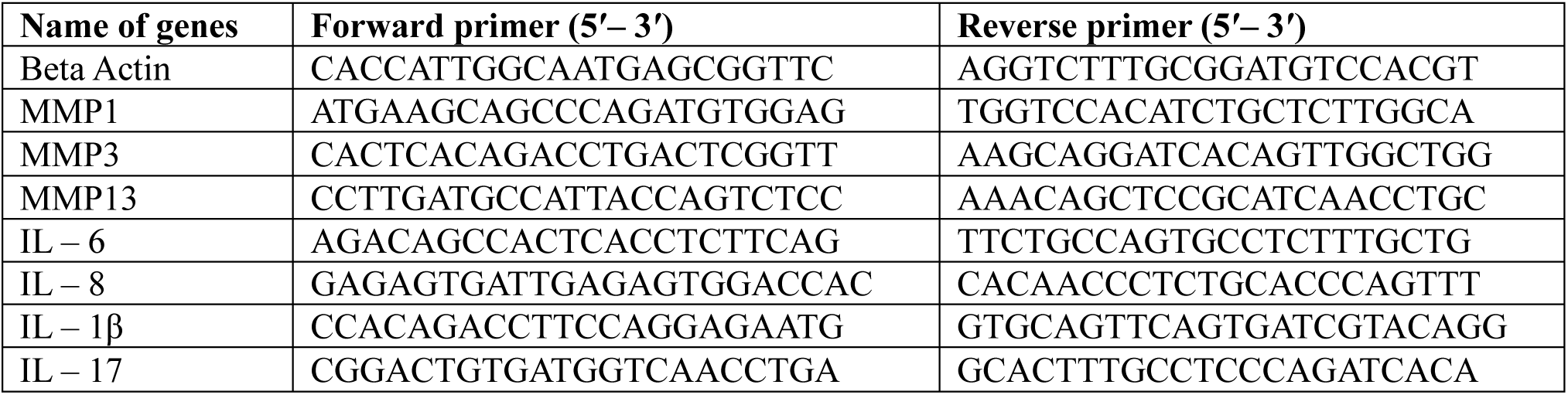
Forward and reverse primer sequences for the inflammatory cytokine genes.

##### 2.2.6.3. Anti-inflammatory effect evaluation

After the selection of the overexpressed cytokines from the cytokine profiling, six different formulations were prepared and tested, along with a positive and a negative control. The cells (C28/I2 and primary chondrocytes) were seeded separately in 6-well plates at a seeding density of about 30,000 cells per well. At around 50% confluency, the cells were treated with 10 ng/mL of IL-1β for 24 hours to create inflammatory conditions, except the negative control. Chondrocytes were transfected one day after seeding with formulations and incubated for another 48 hours. The inflammatory markers were evaluated in both the C28/I2 cell line and the primary chondrocytes after induction of inflammation by IL-1β. The mRNA levels of the cytokines were measured by qRT-PCR as described above in section 2.2.6.2. The protein expression was quantified using ELISA from cells separately seeded into 6-well plates and treated as given in Table 2, except for the negative control cells and the scramble siRNA lipoplexes. At the end of the incubation time, the supernatant cell culture medium was collected, frozen, and stored at −80°C. The concentrations of IL-6 and IL-8 were checked with ELISA kits (Human IL-6 Standard TMB ELISA Development Kit & Human IL-8 (CXCL8) Standard TMB ELISA Development Kit, Peprotech, USA). The ELISA for IL-6 and IL-8 was performed according to the manufacturer’s protocol. The optical density (OD) values were recorded using the plate reader FLUOStar from BMG Labtech (Ortenberg, Germany).

**Table 2.** Conditions/treatments tested on inflamed cells.

| S.No | Name of condition | IL-1β induced | Treatment | Abbreviation |
| --- | --- | --- | --- | --- |
| 1 | Positive control (IL-1β-induced and untreated cells) | ✓ | X | PC |
| 2 | Negative control (non-induced and untreated cells) | X | X | NC |
| 3 | Curcumin-loaded cationic liposomes (optimal formulation) | ✓ | ✓ | Opt-CL |
| 4 | IL-6 and IL-8 siRNA + Curcumin co-loaded lipoplexes | ✓ | ✓ | ILsiRNA-CLx |
| 5 | Scramble siRNA + Curcumin co-loaded lipoplexes. | ✓ | ✓ | SsiRNA-CLx |
| 6 | IL-6 and IL-8 siRNA lipoplexes | ✓ | ✓ | ILsiRNA-Lx |
| 7 | Scramble siRNA lipoplexes | ✓ | ✓ | SsiRNA-Lx |
| 8 | IL-6 and IL-8 siRNA with Lipofectamine RNAimax | ✓ | ✓ | ILsiRNA-LR |

IL-1β-induced cells without treatment were termed positive control (PC), and non-induced cells without treatment were termed negative control (NC). The CL obtained through DoE (Opt-CL), which was devoid of any oligonucleotides, was tested to check the effect of curcumin alone. Besides the optimal formulation, several types of lipoplexes were prepared and tested. To obtain the co-loaded lipoplexes-based treatments, the therapeutic or scramble (negative control) siRNA was added to the Opt-CL in an N/P ratio of 20:1, following the procedure described in 2.2.3.3. Thus, co-loaded lipoplexes obtained through the complexation of Opt-CL with IL-6 and IL-8 siRNA (ILsiRNA-CLx) were used to evaluate any synergistic effects of curcumin and the mixture of the oligonucleotides.

Additionally, CL complexed with scrambled siRNA (SsiRNA-CLx) were used to evaluate off-target effects or nonspecific responses. Parallelly, lipoplexes without curcumin were prepared by complexing liposomes having the same lipid composition and the same physicochemical properties as the Opt-CL but without curcumin, with IL-6 and IL-8 (ILsiRNA-Lx) to evaluate the anti-inflammatory efficacy of siRNA alone, without the influence of curcumin. Scramble siRNA lipoplexes (SsiRNA-Lx) were prepared to observe the off-target effects without curcumin. Finally, Lipofectamine RNAimax with oligonucleotides (ILsiRNA-LR) was taken as a reference for efficacy. The list of the treatments and their abbreviations is given in Table 2.

#### 2.2.7. Assessment of the antioxidant effect

##### 2.2.7.1. Induction of oxidative stress and treatment

The induction of oxidative stress by hydrogen peroxide has been studied extensively on chondrocyte cells by D’Adamo et al. [28]. A similar procedure has been tested and implemented for this study. The C28/I2 chondrocyte cell line and the primary chondrocytes were seeded in T-75 flasks. At around 80% confluency, oxidative stress conditions were induced by adding 0.5 mM hydrogen peroxide, and the cells were further incubated overnight. The following day, cells were washed with PBS, and CL equivalent to 50 µM of curcumin were added directly to the cell culture media. The cells were incubated for an additional 24 hours after transfection, washed with PBS, and lysed using the RIPA lysis buffer. The cell lysates were analyzed for the antioxidant activity of curcumin.

##### 2.2.7.2. Evaluation of oxidative stress markers

The antioxidant activity of curcumin is known to reduce the oxidative stress induced by hydrogen peroxide in chondrocytes [29]. The cell lysates obtained from the previous step were analyzed in terms of total oxidant status (TOS), total antioxidant capacity (TAC), and malondialdehyde (MDA) level. An automated colorimetric method, developed by Erel, was used to measure the TOS in the cell lysates [30]. Samples were prepared according to established protocols and subjected to the procedure previously described by Porfire et al. [20] to conduct the assay. Standard calibration curves converted the colorimetric readings obtained into TOS values. The TOS results were expressed in terms of mmol H_2_O_2_ equivalent/L. The TAC assay, adapted from Erel et al., assesses the capacity of antioxidants within a solution to decolorize the blue-green ABTS+ radical cation (2,2-azinobis(3-ethylbenzothiazoline-6-sulfonate)), which varies according to their concentrations and antioxidant capacities [31]. The procedure for the assessment followed was adapted from previous studies from our group [32,33]. In order to assess lipid peroxidation, a quantitative analysis of MDA was performed, previously reported by Patras et al. [34]. MDA concentration was normalized to the protein concentration in each cell lysate.

### 2.3. Statistical analysis

All measurements are expressed as mean ± standard deviation. The statistical analysis related to the QbD approach was conducted using MODDE 13 Pro software. The primary goal of this analysis was to fit the experimental data into a statistical model, ensuring a robust understanding of the relationships between the variables. A summary of fit and ANOVA were performed to validate the model’s predictive capability, ensuring that variability in the data was adequately explained. Coefficient plots and response contour plots were generated to assess the influence of individual variables on the system’s output. From this analysis, set points were identified to achieve the desired responses, and a design space plot was constructed to explore operational boundaries further.

For the in vitro experimental results, GraphPad Prism version 10.2.2 was used for statistical analysis and graph generation. An Area Under The Curve (AUTC) graph was created to represent curcumin release over time. An unpaired t-test was performed to compare the differences in curcumin release profiles between conditions with pH 5 and pH 6.8 in the presence of 1% Tween-80. The unpaired t-test was chosen because it allows for comparison between two independent groups to determine whether there is a statistically significant difference between their means.

ANOVA was used to compare the means of multiple groups (e.g., control vs. various treatment conditions) to determine whether any significant differences existed between the groups. This method was selected because it enables the analysis of differences across multiple conditions simultaneously, rather than performing multiple t-tests, which could increase the risk of Type I error. The ANOVA results help to validate the reliability of the experimental conditions and their influence on the outcomes.

Statistical significance was defined based on P-values, with results considered significant when P < 0.05 (*P < 0.05; **P < 0.01; ***P < 0.001; ****P < 0.0001). Non-significant differences were reported when P > 0.05 (n.s.). This threshold ensures that any observed differences in the data are unlikely to be due to random variation alone, thus validating the experimental results.

## 3. Results and discussion

### 3.1. QbD approach

#### 3.1.1. Defining the QTPP and risk assessment through FMEA

Cationic liposomes (CL) are considered the most promising delivery vehicles for genetic material due to their adequate biodegradability and biocompatibility. Additionally, their preparation is relatively simple, and for their formulation, cationic lipids allow the binding of nucleic acids through electrostatic interactions. Today, when the complexity of the pathophysiological mechanisms has been deciphered in many diseases, nucleic acids are associated with other molecules in therapy to target as many pathological phenomena as possible. For this reason, developing more complex delivery systems that can successfully co-deliver nucleic acid and the associated active substance in the tissue of interest is needed, and the design of such delivery systems is more complex. In our study, we employed a QbD approach to facilitate the development of CL, which was further complexed with siRNA to get co-loaded lipoplexes. In line with the QbD approach, the QTPP was first established for lipoplexes, along with identifying the CQAs, as detailed in Table 3. According to ICH Q8(R2), a CQA is “a physical, chemical, biological, or microbiological property or characteristic that should be within an appropriate limit, range, or distribution to ensure the desired product quality” [22].

**Table 3.** QTPP for co-loaded lipoplexes intended for intra-articular delivery.

| QTPP aspect | Requirement |
| --- | --- |
| Therapeutic Target | Deliver genetic material (siRNA, mRNA, or plasmid DNA) and curcumin to chondrocytes. Achieve effective gene silencing or overexpression of target genes. Providing therapeutic benefits by reducing inflammation and promoting cartilage regeneration in osteoarthritis. |
| Product characteristics | Ability to complex oligonucleotides and to encapsulate curcumin within stable and biocompatible CL. Average particle diameter suitable for cellular internalization (~100-200 nm) with uniform size distribution. Positive surface charge (to facilitate interaction with negatively charged cell membranes (~30 mV)). Incorporation of pegylated phospholipids and cholesterol to avoid RES clearance and enhance cellular uptake and stability. High encapsulation efficiency for curcumin (>80%) and good complexation capacity for oligonucleotide (>80%). A sustained release profile to ensure a prolonged therapeutic effect. |
| *In vitro* performance | Low cytotoxicity to maintain cell viability (>80%) and minimize adverse effects on chondrocytes. Efficient cellular internalization and transfection (>50% transfection efficiency) in vitro. |
| Stability and Storage | Long-term stability of liposomes under storage conditions (e.g., 4°C) without significant degradation or aggregation. Resistance to environmental factors such as light, temperature fluctuations, and oxidation. Retention of therapeutic activity over extended periods to support shelf-life requirements. |

Based on available scientific literature and pre-formulation studies, the particle size (Y1), PDI (Y2), surface charge (Y3), encapsulation (Y4), and complexation (Y5) capacities, fluorescence levels (Y6) along with cell viability (Y7) and transfection (Y8) were chosen as CQAs of our liposomal vector and were optimized through DoE.

An Ishikawa diagram in Figure S.2 helped us identify the CMAs and CPPs linked to the CQAs of the liposomes. A risk assessment was done on all the crucial variables chalked in the diagram, and the RPN was calculated. As described in this study, six variables with high RPN (> 150) were selected for further investigation by experimental design. The variables identified through risk analysis (Table S.1) via FMEA are discussed in detail in relation to formulation parameters, process parameters involved in the preparation of liposomes and lipoplexes, and the parameters used for *in vitro* study evaluation parameters.

The main variables identified were dependent on the lipid composition, which plays a critical role in the development of liposomes to meet the established QTPP. It significantly affects the liposomal structure, influencing both its physicochemical properties and biological performance. In our study, we have methodically selected a range of lipids, encompassing 1,2-dioleoyl-3-trimethylammonium-propane (DOTAP) as the cationic lipid, and a spectrum of helper lipids: 1,2-dioleoyl-sn-glycero-3-phosphoethanolamine (DOPE), 1,2-dioleoyl-sn-glycero-3-phosphocholine (DOPC), 1,2-dipalmitoyl-sn-glycero-3-phosphocholine (DPPC), methoxy(polyethylene glycol)-distearoylphosphatidylethanolamine (MPEG-DSPE-2000), and cholesterol. Each lipid component was chosen based on its unique attributes and potential synergistic effects within the liposomal matrix. DOTAP, recognized for its cationic nature, is one of the most widely used lipids in cationic liposomal formulations, facilitating efficient cellular uptake and intracellular cargo delivery. Its inherent positive charge facilitates electrostatic interactions with negatively charged cell membranes, enhancing cellular internalization and endosomal escape of encapsulated payloads. Many studies highlight DOTAP’s versatility in nucleic acid delivery and small molecule therapeutics, emphasizing its significance in biomedical applications [35].

The mammalian cell membrane consists of phosphatidylethanolamine and phosphatidylcholine; hence, liposomes containing lipids of this kind could fuse or interact more readily with the membranes, helping cell internalization. Helper lipids such as DOPE, DOPC, and DPPC complement the formulation strategy and help modulate membrane fluidity, stability, and fusogenicity. DOPE, an anionic lipid, augments membrane curvature and promotes endosomal escape, enriching the liposomal delivery efficacy. DOPC, characterized by its zwitterionic nature, enhances biocompatibility and fluidity, contributing to liposomal stability and stealth properties. Conversely, DPPC, a saturated phospholipid, reinforces membrane rigidity and stability, improving encapsulation efficiency and circulation half-life [36].

Furthermore, the incorporation of MPEG-DSPE-2000 and cholesterol further diversifies the lipid matrix, presenting additional functionalities and enhancing liposomal performance. MPEG-DSPE-2000, a polyethylene glycol (PEG)-modified lipid, imparts steric stabilization and stealth properties, helps escape phagocytosis, and reduces immunogenicity. Cholesterol, a crucial component in lipid bilayers, regulates membrane fluidity and permeability, influencing liposomal stability and cargo release kinetics. By methodically investigating the relationship between these diverse lipid components within the QbD framework, we aimed to elucidate their collective impact on cationic liposomal formulation attributes, paving the way for tailored delivery systems with enhanced efficacy and therapeutic potential [37].

As mentioned in the QTPP, particle sizes of less than 200 nm are considered optimal for cellular uptake. The PDI of these liposomes, less than 0.2, would benefit a homogenous population, ideally ensuring that the cellular uptake does not have much variability [38,39]. In this study, the liposomes were extruded through polycarbonate membranes at high pressure (50-75 psi) and a temperature of 45°C, above the transition temperature of the phospholipids. Another critical attribute of the nanosystem that influences cellular uptake and stability is the surface charge of the liposomes. Since the cell membrane is negatively charged, the positive charge of the liposomes facilitates the attachment and delivery of the drug and genetic material. High surface charges on liposomes may induce cytotoxicity, whereas lower surface charges can lead to instability and agglomeration. Hence, an ideal surface charge of around +30 mV was targeted. The cationic lipid concentration is the most critical factor contributing to the charge, although pegylation may reduce the charge.

Due to the complex nature of the developed systems, the curcumin encapsulation in the CL and their ability to complex the oligonucleotides are critical. The EE of curcumin is essential for delivering higher drug concentrations at a lower administered dose. The type of helper lipids, along with their and other phospholipids’ concentrations, were reported to influence the encapsulation of the drug material. The complexation capacity of these liposomes, used as vehicles for the delivery of oligonucleotides, is primarily related by their surface charge. The N/P ratios, representing the molar ratio of nitrogen (from liposomal amines) to phosphate (from genetic material), directly affect the extent of complexation between the liposomes and the genetic material. Therefore, liposomes with higher complexation efficiency are optimal for effective delivery, as they ensure better interaction and encapsulation of the genetic material. [40].

#### 3.1.2. Design of Experiments

The efficient optimization of experimental parameters is fundamental to developing any pharmaceutical product. To achieve this objective, we have utilized the DoE methodology, which offers a systematic approach to study the effects of multiple variables simultaneously. Unlike traditional one-factor-at-a-time (OFAT) approaches, which can be time-consuming and yield suboptimal results, DoE allows for the exploration of complex interactions among factors in a structured and efficient manner. After the risk analysis, we have identified and selected 4 variables, out of which the helper lipid type (X1), the concentrations of cationic lipid (DOTAP) (X2), cholesterol (X3), and pegylated phospholipid (X4), to be investigated by DoE. Among the selected variables, X1 was a qualitative variable, while the rest were quantitative-type variables assessed at three levels: low, medium, and high. The levels of the variables X1-X4 are mentioned in Table 4.

**Table 4.** DoE matrix presenting the variable settings along with the responses.

| Exp Name | X1 | X2 | X3 | X4 | Y1 | Y2 | Y3 | Y4 | Y5 | Y6* | Y7* | Y8 |
| --- | --- | --- | --- | --- | --- | --- | --- | --- | --- | --- | --- | --- |
| N1 | DOPE | 5 | 1 | 1 | 142.20±0.38 | 0.20±0.02 | 38.10±0.47 | 54.12±0.29 | 48.29 | 502.83±56.36 | 102.74±1.67 | 62.46±6.27 |
| N2 | DOPE | 7.5 | 3 | 1 | 159.40±1.06 | 0.14±0.02 | 46.40±0.87 | 91.92±0.49 | 97.35 | 369.83±24.09 | 108.65±0.97 | 73.50±19.13 |
| N3 | DOPE | 10 | 5 | 1 | 161.90±1.16 | 0.13±0.03 | 46.90±0.58 | 90.63±0.89 | 76.83 | 546.50±16.80 | 102.91±2.39 | 55.27±17.72 |
| N4 | DOPC | 5 | 1 | 1 | 176.80±1.30 | 0.10±0.04 | 42.60±4.13 | 92.79±1.13 | 31.74 | 339.50±45.18 | 96.06±2.92 | 49.71±22.86 |
| N5 | DOPC | 7.5 | 3 | 1 | 182.70±1.36 | 0.10±0.03 | 44.90±0.64 | 89.54±1.07 | 100 | 579.83±19.40 | 100.00±6.30 | 58.03±16.42 |
| N6 | DOPC | 10 | 5 | 1 | 156.10±2.86 | 0.18±0.02 | 43.90±0.25 | 85.17±0.06 | 100 | 545.50±88.94 | 109.85±5.10 | 59.48±8.10 |
| N7 | DPPC | 5 | 3 | 1 | 177.80±0.91 | 0.07±0.01 | 45.20±0.21 | 95.89±1.00 | 98.68 | 341.50±8.96 | 101.11±2.98 | 43.68±30.43 |
| N8 | DPPC | 7.5 | 5 | 1 | 182.30±2.10 | 0.08±0.03 | 42.00±1.97 | 90.38±0.77 | 91.59 | 772.17±54.51 | 101.54±1.12 | 41.66±27.47 |
| N9 | DPPC | 10 | 1 | 1 | 177.00±1.25 | 0.12±0.03 | 46.80±1.89 | 96.03±2.05 | 96.16 | 613.83±107.86 | 104.97±1.32 | 52.66±5.74 |
| N10 | DOPE | 5 | 5 | 2 | 170.40±0.81 | 0.11±0.01 | 36.90±1.61 | 96.44±0.71 | 100 | 414.17±112.86 | 99.66±1.32 | 53.75±15.17 |
| N11 | DOPE | 7.5 | 1 | 2 | 150.10±2.66 | 0.16±0.02 | 42.00±2.50 | 87.95±0.85 | 100 | 493.50±34.53 | 107.46±1.03 | 44.79±21.95 |
| N12 | DOPE | 10 | 3 | 2 | 176.50±1.74 | 0.13±0.02 | 45.50±1.18 | 93.49±0.46 | 14.33 | 318.50±15.04 | 103.68±2.06 | 57.43±8.62 |
| N13 | DOPC | 5 | 3 | 2 | 184.50±1.54 | 0.09±0.04 | 33.80±1.14 | 84.02±0.68 | 94.08 | 255.17±15.52 | 90.40±1.55 | 62.33±5.06 |
| N14 | DOPC | 7.5 | 5 | 2 | 175.80±1.75 | 0.07±0.02 | 42.00±0.96 | 85.68±0.41 | 74.13 | 330.50±40.61 | 93.66±2.50 | 60.29±5.09 |
| N15 | DOPC | 10 | 1 | 2 | 180.90±0.96 | 0.06±0.02 | 52.20±0.57 | 83.44±0.91 | 69.73 | 330.83±32.65 | 96.49±3.57 | 42.56±27.53 |
| N16 | DPPC | 5 | 5 | 2 | 176.30±0.68 | 0.10±0.02 | 40.40±0.12 | 94.28±2.46 | 100 | 448.17±69.29 | 111.57±9.83 | 55.03±16.66 |
| N17 | DPPC | 7.5 | 1 | 2 | 173.80±0.32 | 0.10±0.01 | 43.00±0.98 | 90.77±1.85 | 100 | 533.83±64.07 | 105.14±1.61 | 42.98±26.01 |
| N18 | DPPC | 10 | 3 | 2 | 170.10±0.644 | 0.12±0.00 | 42.70±0.75 | 88.43±0.58 | 100 | 225.50±32.02 | 90.92±1.04 | 54.58±17.84 |
| N19 | DOPE | 7.5 | 3 | 1.5 | 139.50±0.55 | 0.16±0.01 | 49.90±1.76 | 91.65±0.47 | 11.17 | 214.50±26.86 | 100.09±3.75 | 64.15±10.78 |
| N20 | DOPE | 7.5 | 3 | 1.5 | 155.80±1.40 | 0.16±0.02 | 42.80±0.42 | 94.11±0.89 | 81.55 | 483.17±16.64 | 105.06±1.75 | 61.05±10.10 |
| N21 | DOPE | 7.5 | 3 | 1.5 | 141.10±0.42 | 0.18±0.02 | 44.20±0.17 | 91.02±0.90 | 100 | 446.50±12.74 | 103.51±3.89 | 62.44±37.13 |

X1 – Helper lipid type, X2 – Concentration of cationic lipid (mM), X3 – Concentration of cholesterol (mM), X4 – Concentration of pegylated phospholipid (mM), Y1 – Liposome size (nm), Y2 – PDI, Y3 – Zeta potential (mV), Y4 – Curcumin encapsulation efficiency (%), Y5 – siRNA complexation capacity (%) for N/P ratio of 2.5:1, Y6 – Fluorescence level, Y7 – Cell viability (%), Y8 – Cell transfection (%)
\*Values corresponding to 50 mM curcumin encapsulated in liposomes

The total lipid content in the liposomal formulation was kept constant at 20 mM. The helper lipid concentration was adjusted according to the concentrations of the remaining lipids. The amount of curcumin taken for encapsulation was also kept constant at 1 mM to evaluate the efficiency among the formulations. The process parameters for all the experiments were maintained constant throughout the design.

Considering the number of degrees of freedom and the power, an L18 (3-level) fractional factorial design was generated with three center points, resulting in 21 experiments. All experiments were performed in a randomized order to eliminate the experiment bias. All the formulations in the DoE were characterized in terms of particle size (Y1), PDI (Y2), zeta potential (Y3), curcumin entrapment efficiency (Y4), siRNA complexation capacity (Y5), fluorescence level (Y6), cell viability (Y7), and cell transfection (Y8). The experimental setup and the responses are represented in Table 4. All experiments were performed in a randomized order to eliminate the experiment bias.

##### 3.1.2.1. Fitting the model and statistical analysis

The experimental data from the responses (Y1 – Y8) were fitted into a multiple linear regression (MLR) model. The statistical parameters that describe the summary of fit are R^2^ (model fit), Q^2^ (future prediction precision), MV (model validity), and MR (model reproducibility). The model parameters for all the responses were estimated and presented in Table 5. According to the literature, values greater than 0.5 for all the parameters indicate a good model. The parameters listed in Table 5 prove that the model fitting is appropriate and indicates a valid and reproducible model. ANOVA and Lack-of-fit tests were also conducted to evaluate the model’s accuracy, demonstrating the significance (p-values < 0.05) and absence of lack-of-fit. The coefficient plots demonstrate the influence of the variables on the studied responses (Figure 1), and the contour plots (Figure 2) were created to depict the predicted response values for two selected variables while keeping the others constant.

**Figure 1.**
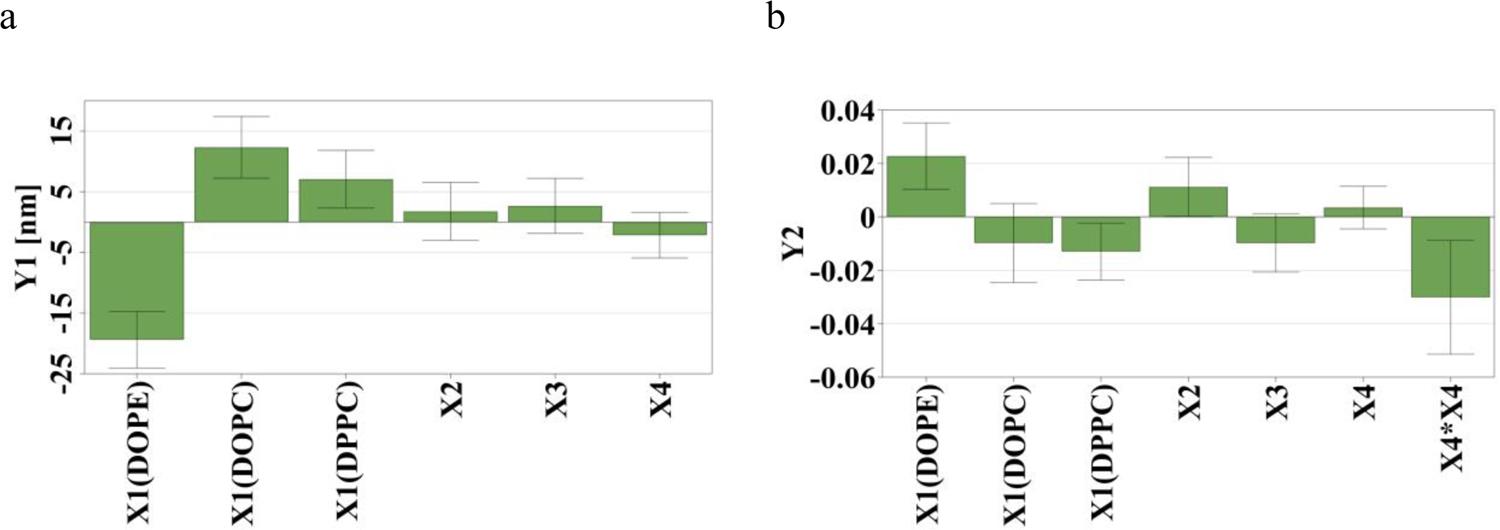
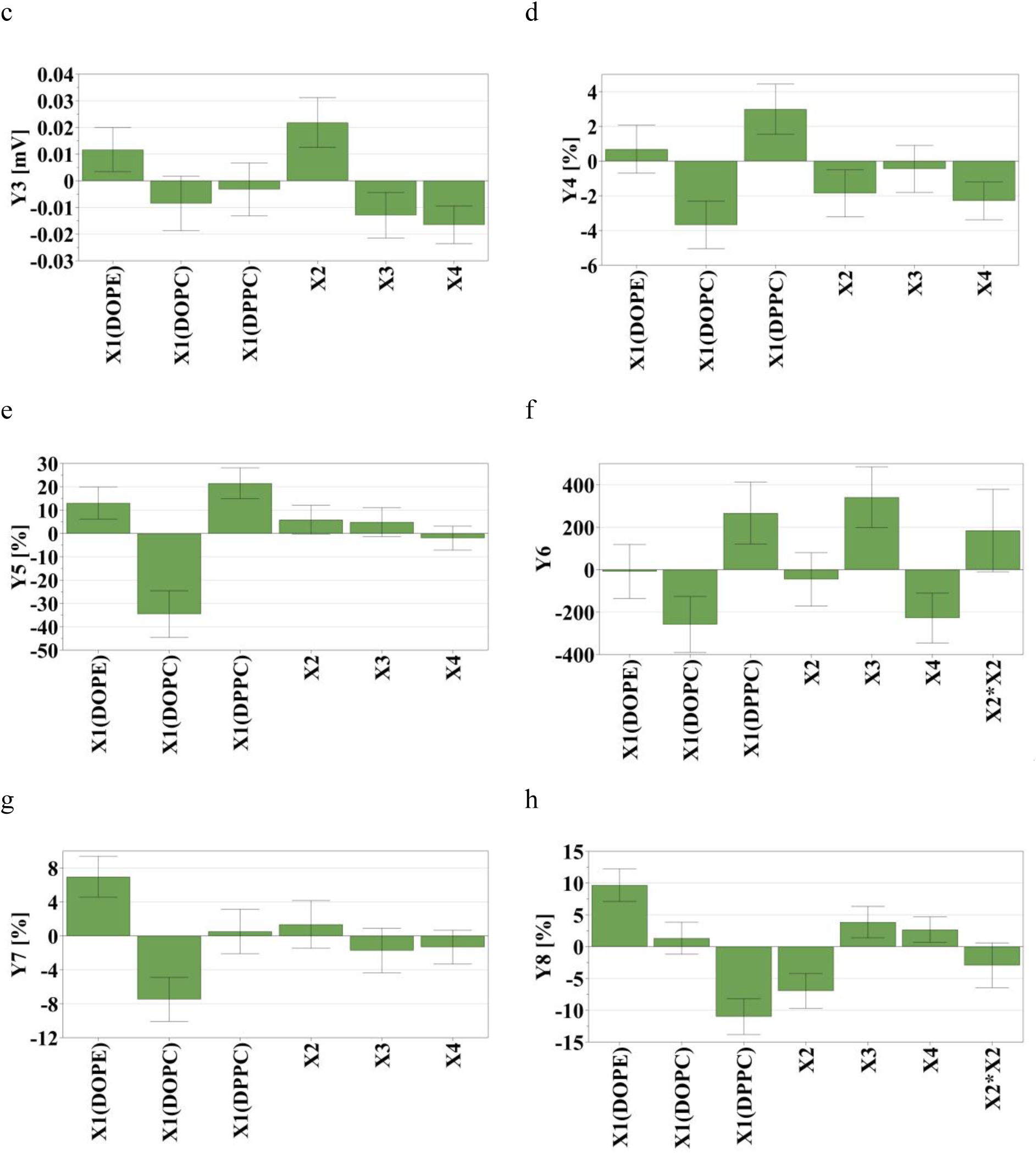
Regression coefficient plots pictorially representing the influence of variables on the responses, a) Particle size (nm) – Y1, b) PDI – Y2, c) Zeta potential (mV) – Y3, d) Curcumin encapsulation efficiency (%) – Y4, e) siRNA complexation capacity (%) – Y5, f) Fluorescence level – Y6, g) Cell viability (%) – Y7, h) Cell transfection (%) – Y8. X1 – Helper lipid type, X2 – Concentration of cationic lipid (mM), X3 – Concentration of cholesterol (mM), X4 – Concentration of pegylated phospholipid (mM)

**Figure 2.**
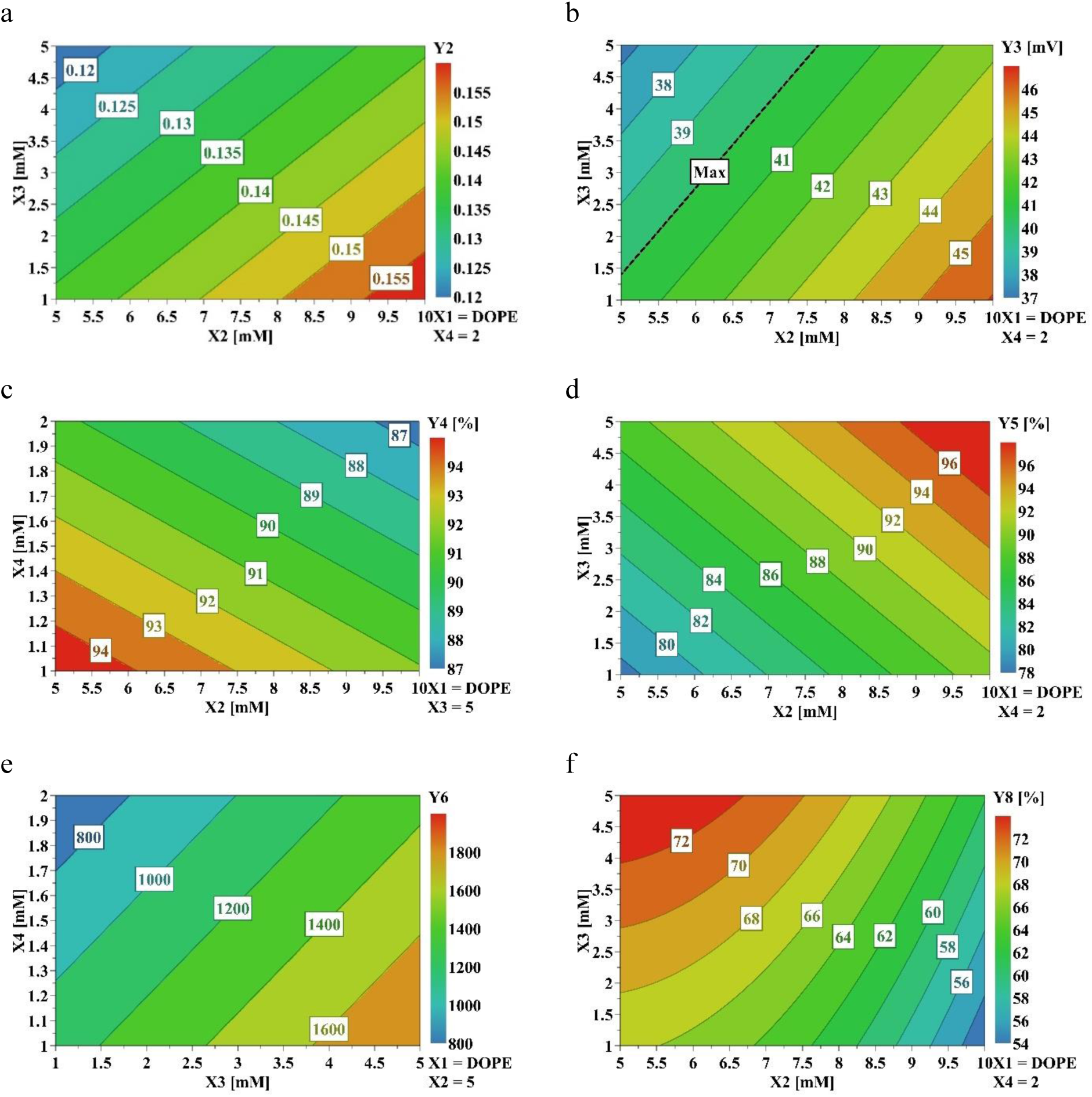
Contour plots showing the interaction between variables and their influence on the responses, keeping DOPE set as a helper lipid for all plots. a) PDI – Y2, b) Zeta potential (mV) – Y3, c) Curcumin encapsulation efficiency (%) – Y4, d) siRNA complexation capacity (%) – Y5, e) Fluorescence levels – Y6, f) Cell transfection (%) – Y8. X1 – Helper lipid type, X2 – Concentration of cationic lipid (mM), X3 – Concentration of cholesterol (mM), X4 – Concentration of pegylated phospholipid (mM).

**Table 5.** Statistical parameters – one-way analysis of variance (ANOVA) test and summary of fit parameters for the studied responses.

| Response | DF | SS | MS (variance) | F - statistic | p-value | Lack of fit | $R^2$ | $Q^2$ | MV | MR |
| --- | --- | --- | --- | --- | --- | --- | --- | --- | --- | --- |
| Y1 | 12 | 501.05 | 41.754 | 16.995 | 0.000 | 0.857 | 0.876 | 0.761 | 0.961 | 0.661 |
| Y2 | 10 | 0.001 | 0.000 | 13.763 | 0.000 | 0.354 | 0.892 | 0.7 | 0.74 | 0.911 |
| Y3 | 9 | 0.001 | 0.000 | 17.205 | 0.000 | 0.518 | 0.905 | 0.731 | 0.836 | 0.915 |
| Y4 | 11 | 36.582 | 3.326 | 12.806 | 0.000 | 0.507 | 0.853 | 0.619 | 0.83 | 0.829 |
| Y5 | 10 | 557.529 | 55.753 | 20.888 | 0.000 | 0.768 | 0.913 | 0.814 | 0.934 | 0.799 |
| Y6 | 9 | 229831 | 25536.8 | 12.148 | 0.001 | 0.163 | 0.89 | 0.645 | 0.545 | 0.991 |
| Y7 | 11 | 115.363 | 10.488 | 12.912 | 0.000 | 0.621 | 0.854 | 0.655 | 0.881 | 0.774 |
| Y8 | 8 | 65.432 | 8.179 | 18.211 | 0.000 | 0.205 | 0.932 | 0.752 | 0.603 | 0.965 |

Y1 – Liposome size (nm), Y2 – PDI, Y3 – Zeta potential (mV), Y4 – Curcumin encapsulation efficiency (%), Y5 – siRNA complexation capacity (%), Y6 – Fluorescence level, Y7 – Cell viability (%), Y8 – Cell transfection (%), DF – Degrees of Freedom, SS – Sum of squares, MS – Mean squares, $R^2$ – Model fit, $Q^2$ – Future prediction precision, MV – Model validity, MR – Model reproducibility

##### 3.1.2.2. Influence of variables on the CQAs of the CL

###### Particle size, polydispersity index, and Zeta potential

The PS, PDI, and ZP are essential physicochemical properties that influence the *in vitro* and *in vivo* behavior of the CL. The size of liposomes is a critical factor in affecting cellular uptake. For chondrocytes, smaller liposomes (typically < 200 nm) are preferred due to their ability to penetrate the dense extracellular matrix of the cartilage more efficiently. Smaller particles have a higher surface area-to-volume ratio, which enhances their interaction with cell membranes and facilitates endocytosis. More giant liposomes may be engulfed less readily by chondrocytes and could be trapped within the extracellular matrix, reducing their uptake efficiency. Interestingly, larger liposomes are also used to lubricate the synovial joints effectively, as described by Sivan et al. [41]. However, smaller PS appears to have an advantage for cellular uptake. In our study, all liposomal formulations were below 200 nm, ranging from 139.5 nm to 184.5 nm. The PDI indicates the uniformity of liposome size distribution [42]. According to our results, the liposomes had a uniform size distribution, where PDI varied between 0.06 and 0.2. The ZP reflects the surface charge of the liposomes and influences their stability and interaction with cells. Nanoparticles could aggregate due to insufficient electrostatic repulsion between them, and generally, a higher surface charge (< −30 mV or > +30 mV) is considered beneficial to overcome aggregation. Since cells such as chondrocytes typically have glycocalyx that is negatively charged, positively charged liposomes (cationic) exhibit enhanced cellular uptake, presumably by binding with negatively charged glycosaminoglycans such as heparan sulfate on the cell surface [43]. However, excessively high positive charges (> +30 mV) could lead to cytotoxicity and non-specific interactions with negatively charged components in the extracellular matrix. A surface charge on the liposomes that could balance the stability and the cell viability is crucial for cellular uptake [44]. As shown in Table 4, all liposomal formulations exhibited a positive surface charge, ranging between 33.8 mV and 52.2 mV, due to the cationic lipid DOTAP.

The coefficient plot in Figure 1 (a) indicates that the only factor that significantly impacted the PS (Y1) was the helper lipid type (X1). Thus, liposomes made from DOPE as helper lipids had significantly lower PS when compared to liposomes containing DOPC or DPPC, probably because ethanolamine in DOPE has a smaller head group than the choline groups in DPPC and DOPC. This smaller head group leads to tighter packing and reduced steric hindrance among lipid molecules, which can result in smaller and more stable liposomal particles. In contrast, the larger choline head groups in DPPC and DOPC can cause more steric hindrance, leading to larger PS [45].

The PDI values were found to be good for all the formulations in the range of 0.07 to 0.2. However, the PDI (Y2) of the liposomes made from DOPE was slightly higher than for those with DOPC or DPPC, as shown in the Figure 1 (b). The liposomes made from DOPC and DPPC might have had more rigid bilayers than those of DOPE, which might be a reason for their lower PDI when extruded [46,47]. More heterogeneous populations of liposomes were obtained when the concentration of cationic lipids (X2) was increased [48]. The opposite was observed when the concentration of cholesterol (X3) was increased. The effect of the pegylated phospholipids (X4) on the uniformity of the liposomes seemed insignificant; however, the squared factor X4*X4 was inversely proportional to the PDI. For DOPE-containing liposomes, a high cholesterol concentration and a low cationic lipid concentration appeared to offer the best PDI, similar to previous observations, as represented in contour plot in Figure 2(a).

The liposomes made from DOPE had the highest ZP (Y3) compared to liposomes formulated with DOPC and DPPC. The concentration of cationic lipids (X2) was the most significant factor positively influencing the ZP, as shown in Figure 1(c). However, although the pegylated phospholipid and cholesterol are electrically neutral, their concentrations (X3 & X4) were inversely correlated to the ZP. The value of ZP was above 40 mV when higher concentrations of cationic lipids were used., which might lead to cytotoxicity. High cationic lipid concentrations and low cholesterol levels apparently gave the highest ZP for DOPE liposomes, as shown in Figure 2 (b). Therefore, to achieve the target ZP of approximately +30 mV, as specified in the QTPP, to minimize cytotoxicity, lower concentrations of cationic lipids and higher concentrations of cholesterol have to be used as represented in the contour plot Figure 2 (b).

To check how the CQAs of cationic liposomes are influenced by the complexation with oligonucleotides, preliminary studies were done to evaluate the physicochemical properties of lipoplexes co-loaded with curcumin and luciferase siRNA at N/P ratios of 2.5:1, 5:1, 10:1 and 20:1. It was observed that the complexation did not cause significant PS and PDI changes in lipoplexes when compared to cationic liposomes. On the other hand, the ZP (Y3) was influenced by the N/P ratio. Thus, the ZP increased with the increase in the N/P ratios, which denotes that at higher N/P ratios (20:1), there are a higher number of free positive charges on the lipoplex. On the contrary, the ZP decreased at lower N/P ratios (2.5:1 and 5:1), indicating the nucleic acid masking of the surface charge of the lipoplex.

###### Encapsulation efficiency and complexation capacity

The EE (Y4) of curcumin and the CC (Y5) with the oligonucleotides are CQAs of the delivery system that could influence the dosage. Improving these responses means having a higher payload, which requires a lower dose for the same intended effect.

The EE of curcumin was found in the range of 54.12% to 96.44%, whereas the CC for the liposomes at N/P ratio of 2.5:1 ranged broadly from 14.33% to 100%. As seen in Figure 1 (d), the EE (Y4) was high for the liposomes with DPPC and, to a lesser extent, DOPE as helper lipids. On the other hand, using DOPC led to less curcumin being entrapped in the liposomes. The cationic lipid (X2) and pegylated phospholipid (X4) concentrations have been found to impact the EE significantly and inversely. In our study, the cholesterol concentration had an insignificant effect on the EE of curcumin; however, it is known to be directly proportional to the EE [24]. Hence, contour plots were generated using constant values of cholesterol (5 mM) and DOPE as helper lipids. In the contour plot in Figure 2 (c), it was observed that the highest EE was found when low concentrations of cationic lipid and pegylated phospholipid were used.

The complexation capacity (CC) of lipoplexes formed between cationic liposomes (CL) of all 21 trials and luciferase siRNA were evaluated using gel electrophoresis at N/P ratios of 2.5:1, 5:1, 10:1 and 20:1, as depicted in Figure S.3. CC was quantified by measuring the intensity of uncomplexed nucleic acid bands at each ratio of the lipoplex. It was found that at higher N/P ratios greater than 2.5:1, most of the lipoplexes have complete complexation. Hence, in order to differentiate and evaluate the impact of the factors, the CC was calculated at the lowest N/P ratio which was 2.5:1 and the data was summarized in Table 4. Similar to curcumin EE, the liposomes composed of DOPE and DPPC proved to be superior when compared to that of the DOPC liposomes, as shown in Figure 1 (e) mirroring the trend observed in curcumin encapsulation efficiency (EE). Based on these findings and from the previous responses, DOPE was selected as the preferred helper lipid for further studies. Among the various N/P ratios tested, DOPE liposomes achieved complete complexation at a 20:1 ratio. While the concentration of cationic lipids (X2) and cholesterol level (X3) directly influenced CC, their effects were not statistically significant. As illustrated in Figure 2 (d), optimal complexation (over 95 %) was achieved with DOPE as the helper lipid, 2 mM pegylated phospholipid, and high concentrations of both cationic lipid and cholesterol. These results provide valuable insights into the formulation parameters that enhance siRNA complexation in liposomal systems.

###### Cell internalization, cell viability, and cell transfection

The co-loaded lipoplexes are desired to have high cell compatibility, internalization, and therapeutic outcomes. Curcumin is a fluorescent compound that, when internalized into cells, still emits fluorescence, which can be detected by microscopy (Figure S.4) and fluorescence plate readers. This way, using curcumin as a model drug has also helped us estimate cell internalization by measuring the fluorescence intensity from the CL that have passed through the cell membrane. In preliminary studies, the fluorescence of all the formulations was recorded at adjusted curcumin concentrations of 25-100 μM. However, in the DoE, only the fluorescence level (Y6) and cell viability (Y7) recorded at a concentration of 50 μM curcumin were considered, as free curcumin (taken as standard) appeared to be cytotoxic beyond this level. Cell transfection (Y8), on the other hand, represented the gene expression/silencing within the cells. It was studied with the highest N/P ratio (20:1), where it was observed that all the genetic material is bound to the liposomes (Figure S.3). Comparing the 21 DoE formulations allowed to evaluate the release of genetic material after internalization and crossing the cell membrane.

The coefficient plot in Figure 1 (f) reveals that DOPE was insignificant for the fluorescence levels (Y6) amongst the helper lipids, whereas DPPC liposomes were observed to have the highest fluorescence, and the liposomes with DOPC had the lowest. The cationic lipid concentration (X2) along with its non-linear factor X2*X2, are insignificant. However, X2*X2 had a directly proportional relationship with cell internalization. Higher cholesterol levels (X3) seemed to facilitate the localization of the liposomes into the cells; however, the pegylated phospholipid concentration (X4) appeared to have decreased the level of liposome internalization. Considering DOPE as the helper lipid and cationic lipid concentration (X2) at 5 mM for contour plots in Figure 2 (e), we observe that the fluorescence levels are highest over 4 mM of cholesterol (X3) and low levels of pegylated phospholipid (X4).

When the formulations were evaluated regarding the impact on cell viability (Y7), the results showed that the use of DOPE granted the highest cell viability, whereas DOPC use was inversely linked to this response, and the DPPC effect was insignificant as shown in the Figure 1 (g). The other formulation variables, i.e., the concentrations of cationic lipid (X2), cholesterol (X3), and pegylated phospholipid (X4), were also observed to have an insignificant impact on the viability of chondrocytes.

The results were similar for the cell transfection (Y8), showing that the liposomes with DOPE had a higher gene knockdown for the luciferase activity compared to DOPC and DPPC, as shown in Figure 1 (h). Higher concentrations of cationic lipid (X2) were observed to have shown a negative effect on gene knockdown. Cell transfection was more effective with liposomes containing less cationic lipid (X2). However, the cholesterol (X3) and pegylated lipid (X4) concentrations significantly and directly affected cell transfection. At low levels of cationic lipid (X2) and high levels of cholesterol (X3), response contour plots in Figure 2 (f) show high cell viability (Y7) and high cell transfection (Y8).

##### 3.1.2.3. Design space and validation

The design space is a graphical representation of a multidimensional space of operating ranges within which the CQAs of a product are expected to be met. In this study, the design space represented by green in Figure 3 describes the range of the variables that achieve desired response levels for the PS, PDI, ZP, EE, cell viability, and cell transfection with the least probability of failure. The complexation capacity of N/P ratios higher than 10:1 was 100%. Since the amount of cationic liposomes taken at higher ratios did not induce cytotoxicity, a 20:1 N/P ratio was considered to ensure complete complexation, confirmed by the gel electrophoresis method (Figure S.5). Hence, the complexation capacity was not considered as a response for design space exploration.

**Figure 3.**
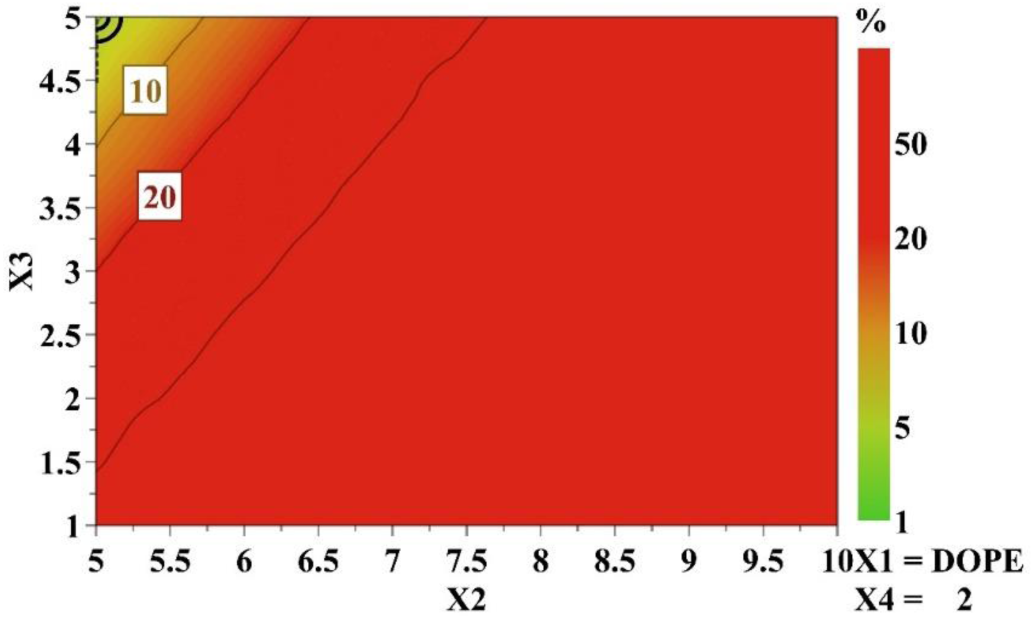
Design space for the CL represented as a function of cationic lipid concentration (X2) and cholesterol concentration (X3) for a constant value of 2 mM pegylated phospholipid concentration (X4) and DOPE (X1) as helper lipid

**Figure 4.**
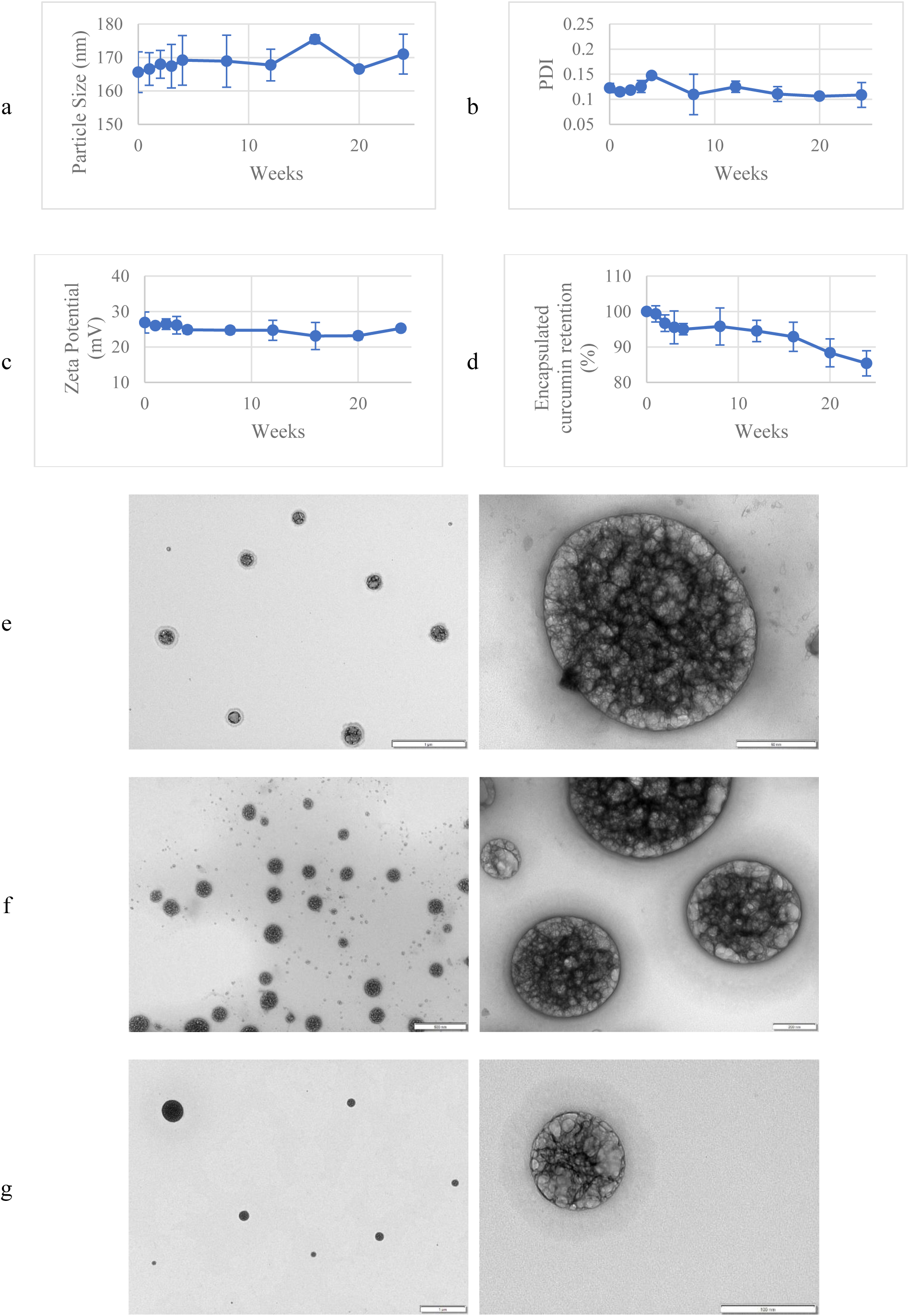
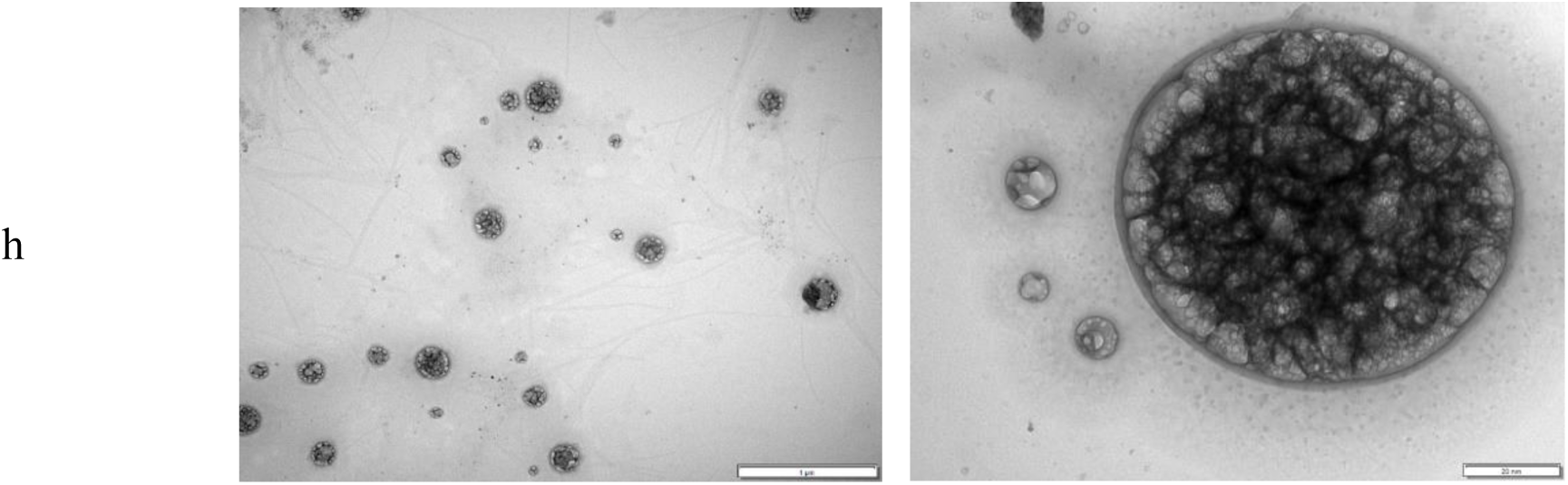
Physicochemical characterization of the freshly prepared Opt-CL over 24 weeks: a) Particle Size, b) Polydispersity Index, c) Zeta potential, d) Encapsulated curcumin retention (%). Results are reported as mean ± standard deviation of three measurements. TEM images of the liposomes: e) Blank liposomes before lyophilization, f) Opt-CL before lyophilization, g) Blank liposomes after lyophilization, h) Opt-CL after lyophilization.

**Figure 5.**
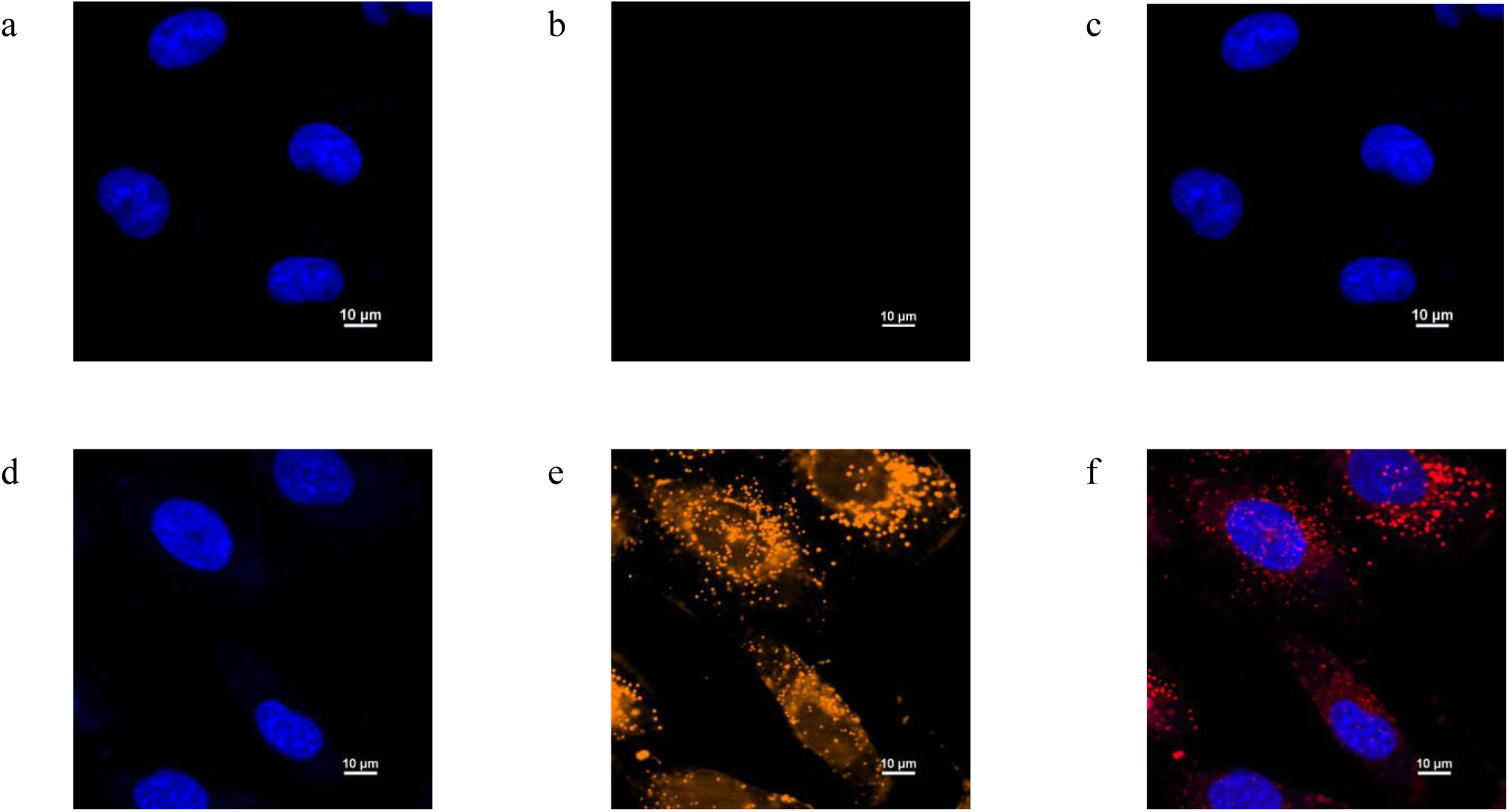
Confocal microscopy images of the control C28/I2 cells treated with Cy5-labelled siRNA (top – a,b,c) and C28/I2 cells treated with lipoplexes complexed with Cy5-labelled siRNA (bottom – d,e,f). The blue color represents the DAPI-stained nucleus, and the yellow and red colors represent the Cy5-labelled lipoplexes. Images corresponded to 405 nm (RCM 405 – a,d), 561 nm (RCM 561 – b,e), and merged (c,f) pictures.

Based on these results, set points that were likely to achieve the desired responses were identified. Of the 4 variables investigated, DOPE was the helper lipid, and the concentration of pegylated phospholipid was fixed at 2 mM as recommended by the software. The remaining variables, namely the concentrations of cationic lipid (X2) and cholesterol (X3), were plotted to indicate the best levels to achieve the desired CQAs for the liposomes with the least probability of failure, pictorially represented in Figure 3. According to the graphical representation, using lower levels of cationic lipids and higher cholesterol concentrations will ensure the obtaining of CL that meet the established quality profile.

### 3.2. Evaluation of Opt-CL

Based on the design space and the set points available, an optimum formulation of liposomes containing 8 mM DOPE, 2 mM MPEG-DSPE, 5 mM DOTAP, and 5 mM cholesterol was prepared and subsequently characterized for its CQAs. The predicted and observed responses were evaluated for the optimum run and listed in Table 6. Furthermore, gel electrophoresis was conducted to determine the N/P ratio at which the nucleic acid complexation with the liposomes is high. Complete complexation was observed at an N/P ratio of 20:1, as visualized in Figure S. 5.

**Table 6.**
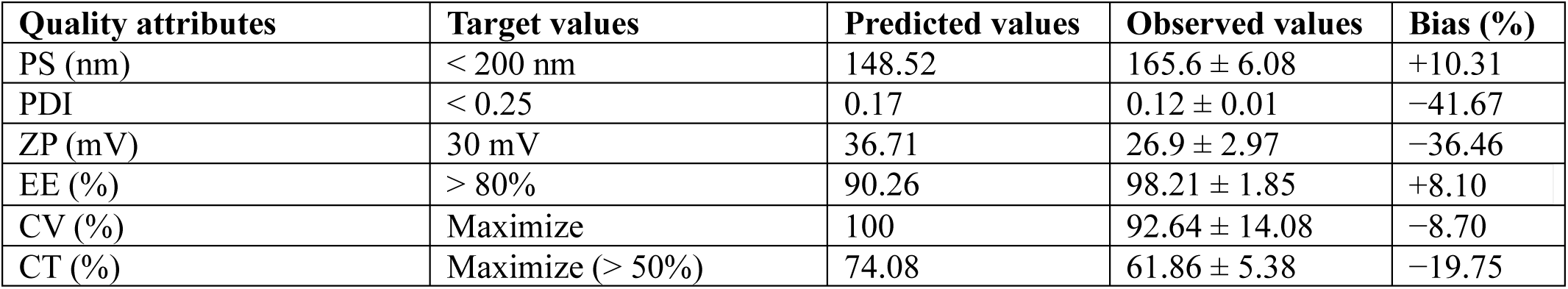
The quality attributes of the Opt-CL formulation: target, predicted, and observed values.

| Quality attributes | Target values | Predicted values | Observed values | Bias (%) |
| --- | --- | --- | --- | --- |
| PS (nm) | < 200 nm | 148.52 | 165.6 ± 6.08 | +10.31 |
| PDI | < 0.25 | 0.17 | 0.12 ± 0.01 | -41.67 |
| ZP (mV) | 30 mV | 36.71 | 26.9 ± 2.97 | -36.46 |
| EE (%) | > 80% | 90.26 | 98.21 ± 1.85 | +8.10 |
| CV (%) | Maximize | 100 | 92.64 ± 14.08 | -8.70 |
| CT (%) | Maximize (> 50%) | 74.08 | 61.86 ± 5.38 | -19.75 |

The Opt-CL obtained through the design space has been further taken forward to demonstrate robustness and efficiency. An optimum formulation with curcumin and a blank formulation with the same lipidic composition as the Opt-CL but without curcumin were prepared. A stability study for the Opt-CL was performed to understand the shelf life of the liposomes in refrigerated conditions. Additionally, the Opt-CL were lyophilized, and their physicochemical properties and morphological rigidity were analyzed. These formulations were then utilized to prove cellular internalization and transfection using confocal microscopy. A release profile was also established to understand the behavior of the liposomes in vivo, in both the inflamed site (pH 6.8) and inside the chondrocytes (lysosomal pH 5).

#### 3.2.1. Stability

One of the primary objectives of this study was to develop physicochemical stable CL to be used as vectors for nucleic acids, so it is crucial to determine the feasibility of long-term storage for the practical application of these formulations. Hence, stability in terms of their physicochemical properties was checked over time at refrigerated conditions. For extended stability, lyophilization was performed, and the reconstituted formulations were also evaluated. Specifically, the study focused on evaluating physical stability, indicated by changes in PS, PDI, ZP, and ECR, as well as morphological stability.

The Opt-CL was regularly tested for six months in terms of PS, PDI, ZP, and the ECR of the liposomes, as represented in Figure 4 a,b,c, and d, respectively. The refrigerated liposomes did not have any significant differences in the physicochemical properties. The PS was 171 nm compared to the initial 165.6 nm, the PDI was observed to be 0.11, and the ZP slightly decreased to 25.25 mV from an initial value of 26 mV. However, the curcumin content was reduced to about 83% by the end of 6 months in refrigerated samples when compared to freshly prepared samples.

For lyophilization, Opt-CL formulated with a 1:1 ratio of trehalose (50 mM) and mannitol (50 mM) as cryoprotectants exhibited optimal PS, PDI, ZP, and curcumin content, closely resembling the initial samples. The stability of both the blank liposomes and Opt-CL was assessed in conjunction with lyophilized and reconstituted liposomes. The initial and final PS, PDI, ZP, and ECR were tested and listed in Table 7. Furthermore, TEM images display their morphology (Figure 4 (e-h)), where it was observed that there were no significant changes in the morphology or physicochemical properties of the liposomes after freeze-drying. The PS has little to no differences, while the PDI has increased from 0.093 to 0.261 for blank liposomes and 0.085 to 0.226 for CL. The ZP also has increased slightly from 24.8 to 30.3 in blank liposomes and from 25.6 to 29.2 in CL. These changes, however, are still within the range of our QTPP and would not affect the safety and efficacy of the liposomal product. Overall, the liposomes maintained much of their initial characteristics, indicating good stability post-lyophilization. Since there were no significant differences between the lyophilized and refrigerated products, either process is considered suitable for the final product.

**Table 7.** Physicochemical properties of the Blank Liposomes and Opt-CL before and after lyophilization.

| Parameters | Before lyophilization |  | After lyophilization |  |
| --- | --- | --- | --- | --- |
|  | Blank liposomes | Opt-CL | Blank liposomes | Opt-CL |
| PS (nm) | 167.0 ± 0.92 | 170.7 ± 0.51 | 171.9 ± 2.17 | 160.1 ± 2.43 |
| PDI | 0.093 ± 0.02 | 0.085 ± 0.02 | 0.261 ± 0.014 | 0.226 ± 0.017 |
| ZP (mV) | 24.8 ± 1.05 | 25.6 ± 0.31 | 30.3 ± 0.40 | 29.2 ± 0.50 |
| ECR (%) | - | 92.34 ± 0.12 | - | 83.14 ± 0.35 |

#### 3.2.2. *In vitro* cell uptake

Cell uptake of the vector is mandatory for the gene silencing efficiency of siRNA cargo. Cellular internalization of the optimum CL was successfully confirmed using fluorescence confocal microscopy, showing that the lipoplexes loaded with the Cy5-labelled siRNA passed through the cell membrane and co-localized with intra-cellular components in the chondrocytes after 24 hours of incubation. Specifically, Figure 10 illustrates a representative fluorescence image after the nuclear staining with DAPI (blue) and after incubation with lipoplexes (red). The uptake of a large number of lipoplexes can be observed in C28/I2 cells, confirmed by the red fluorescence spots inside the living cells.

#### 3.2.3. Release profile

The release studies for the curcumin-loaded liposomes were performed by using the Franz diffusion cells as described by Siefan et al., which were the closest replicable instruments for knee intra-articular space with a 10 mL receptor chamber and liposomes-filled donor chamber divided by the 10 kDa membrane [49]. The sink conditions were ensured, and the release profile for curcumin (Figure 6 a) was performed at pH 5, mimicking the late endosomal pH and pH 6.8, representing the inflamed joint environment [50,51]. Tween 80 was added as a solubilizer to the aqueous media to aid the release of lipophilic curcumin and also gives insight into mimicking the environment in the presence of factors that break down the liposomes [52–54]. A statistical analysis using a t-test was performed after calculating the area under the curve (AUTC) for the release profile in different media (Figure 6 b). Rayamajhi et al. evaluated the endosomal release of doxorubicin-loaded liposomes in this way and observed that within 50 hours, 100% release was achieved at pH 5.5 and about 60% release at pH 7.4 [55]. In our study, the liposomes were stable for more than 72 hours and did not release more than 2% of the drug content at either pH, suggesting no degradation at the injection site or by endosomes. Meanwhile, it was observed that there was a significant increase in the curcumin release of up to 10% in 72 hours at pH 6.8 and up to 15% at pH 5, which denotes a sustained release of curcumin at the injection site.

**Figure 6.**
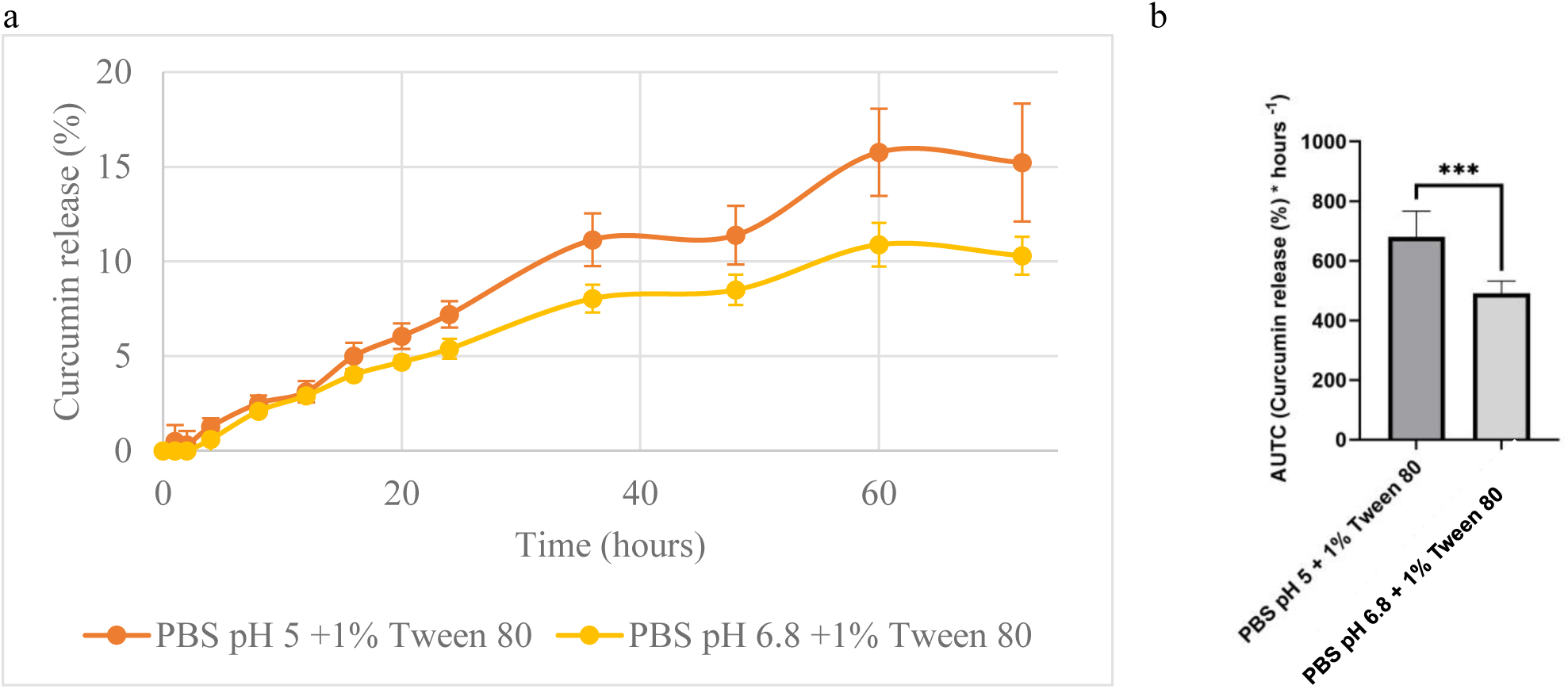
**a)** Release profile data of curcumin from the liposomes in release media at pH 5 and 6.8, with 1% Tween 80, b)Area under the curve (AUTC) graph for the curcumin release from the liposomes in release media at pH 5 and 6.8, with 1% Tween 80. A t-test was performed to analyze the differences between the release media (**, P < 0.01; ***, P < 0.001). The graphs were generated using GraphPad Prism 10.2.2.

The release studies demonstrated that the Opt-CL formulation provided a prolonged release of curcumin, reaching up to 10% in 72 hours at pH 6.8 (representative of inflamed tissue conditions) and up to 15% at pH 5 (endosomal pH). These findings are promising as they suggest a slow, regulated release of the encapsulated curcumin, avoiding an initial burst release at the injection site. This release profile enhances the likelihood of liposome uptake by cells, allowing for intracellular release of curcumin under endosomal conditions, thereby potentially increasing its therapeutic efficacy. The results support the formulation’s suitability for targeted drug delivery, optimizing curcumin bioavailability and effectiveness in the inflamed intra-articular environment. Rayamajhi et al. evaluated the endosomal release of doxorubicin-loaded liposomes in this way and observed that within 50 hours, 100% release was achieved at pH 5.5 and about 60% release at pH 7.4 [55]. While the Opt-CL formulation exhibits a slower release, this is advantageous as it prolongs drug availability and potentially enhances intracellular uptake, making it well-suited for sustained therapeutic action in chronic OA conditions.

### 3.3. Anti-inflammatory effects of the lipoplexes on human chondrocytes

The most significant inflammatory markers found in OA, i.e., IL-1β, IL-6, IL-8, IL-17, MMP-1, MMP-3, and MMP-13, were screened after induction of inflammation with IL-1β in chondrocytes. The mRNA expression levels of proinflammatory cytokines were quantified using qRT-PCR. All the markers were normalized to β-actin for the C28/I2 cell line and the primary chondrocytes (Figure S.6). Through cytokine profiling of inflamed chondrocytes, we found significant overexpression patterns of key inflammatory gene mediators, notably interleukin-6 (IL-6) and interleukin-8 (IL-8), amidst various pro-inflammatory cytokines and matrix metalloproteinases (MMPs) (Figure.S.6) [56]. IL-6 and IL-8 were significantly overexpressed, about 4 and 6 times the relative fold change to the β-actin, in both the IL-1β-induced C28/I2 cell line and the primary chondrocytes, as shown in the supplementary Figure S.6. These observations were similar to those of Kaneko et al., who reported overexpression of IL-6 and IL-8 in the serum and synovial fluid in OA patients [57]. Hence, IL-6 and IL-8 specific siRNA sequences were complexed with the liposomes to yield a delivery system capable of knocking down target genes encoding proinflammatory cytokines. This strategy also allowed us to check to what extent curcumin provides additional benefits to the oligonucleotides in reducing the inflammatory response in chondrocytes.

Figure 7 presents the qRT-PCR and ELISA results representing the mRNA and protein levels of the cytokines in the C28/I2 cell line. The graphs in Figure 7 show that the positive control, representing cells induced by IL-1β, exhibited a significantly higher relative fold change in mRNA levels of both IL-6 and IL-8, as well as elevated protein levels compared to the negative control, which represents non-induced cells, for both cytokines.

**Figure 7.**
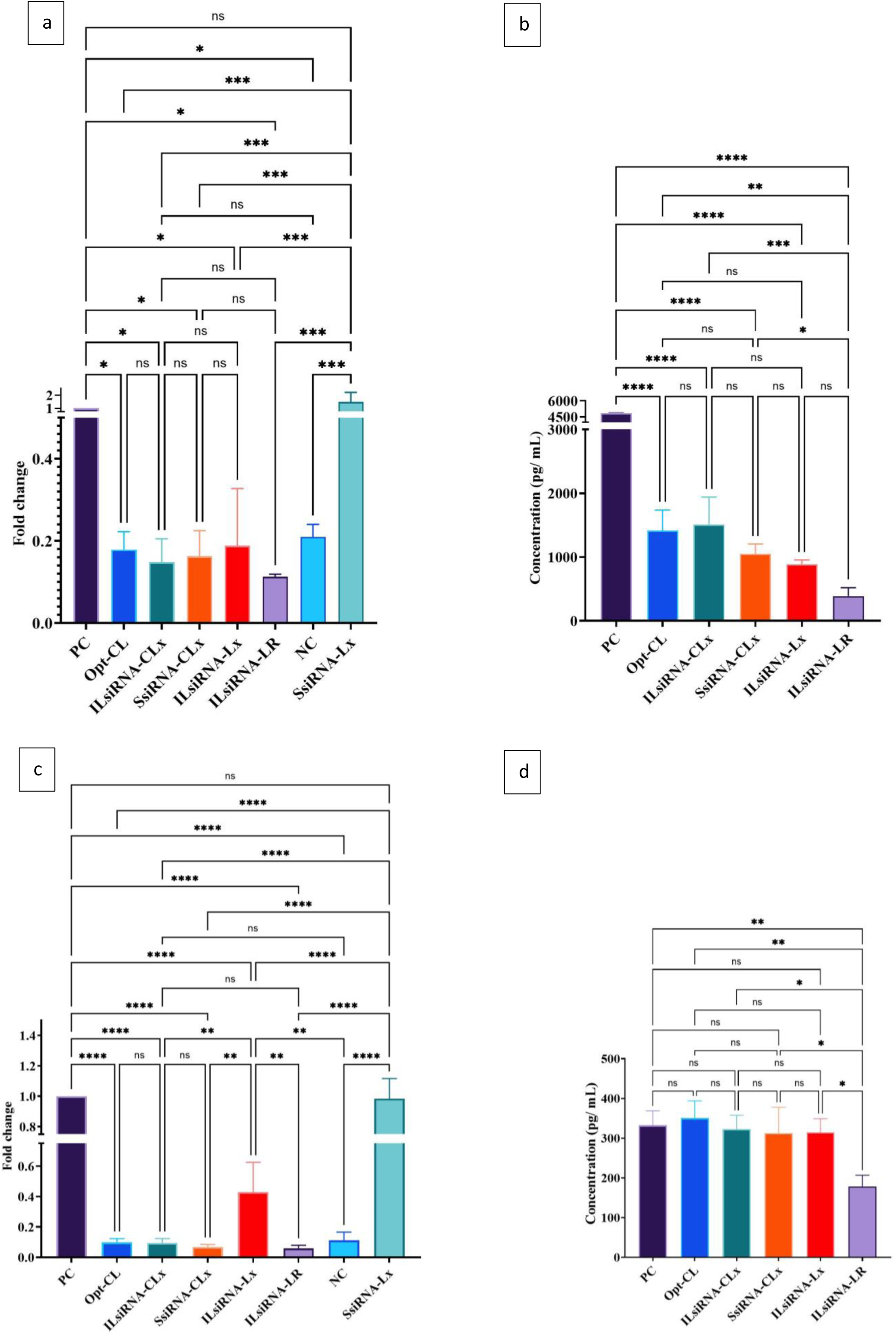
(a) IL-6 mRNA levels in the C28/I2 cell line were expressed through qRT-PCR, and (b) protein levels were quantified using ELISA. (c) IL-8 mRNA levels in the C28/I2 cell line were expressed through qRT-PCR, and (d) protein levels were quantified using ELISA. Positive control (PC), negative control (NC), curcumin-loaded cationic liposomes (Opt-CL), IL-6 and IL-8 siRNA + Curcumin co-loaded lipoplexes (ILsiRNA-CLx), Scramble siRNA + Curcumin co-loaded lipoplexes (SsiRNA-CLx), IL-6 and IL-8 siRNA lipoplexes (ILsiRNA-Lx), Scramble siRNA lipoplexes (SsiRNA-Lx), and IL-6 and IL-8 siRNA with Lipofectamine RNAimax formulation as a standard for transfection and knockdown (ILsiRNA-LR). Results were expressed as mean ± SD of three independent measurements. A one-way ANOVA test with multiple comparisons was performed to analyze the differences between the effects of the treatments (ns, P > 0.05; *, P < 0.05; **, P < 0.01; ***, P < 0.001; ****, P < 0.0001). The graphs were generated using GraphPad Prism 10.2.2.

All the formulations, either liposomes or lipoplexes, induced a significant reduction in the fold change of IL-6 of about 80%, which was almost similar to the uninduced cells (NC). The scramble siRNA lipoplexes (SsiRNA-Lx) proved to have no off-target effects, as they did not have any significant impact on the relative fold change compared to that of the positive control cells. According to Figure 7 (c), the mRNA levels of the IL-8 also showed about an 80% decrease in the fold change for the formulations with curcumin, highlighting its efficacy. The ILsiRNA-Lx, however, was not as effective as the other formulations with Curcumin, with a fold change of about 0.43. The difference in efficacy may be due to curcumin’s broader anti-inflammatory impact, particularly its ability to inhibit pathways that regulate IL-8 expression. Curcumin suppresses both the NF-κB and MAPK pathways, which are key regulators of IL-8 mRNA levels, thereby reducing IL-8 production and its associated inflammatory effects. While the gene silencing strategy employed by ILsiRNA-Lx is also effective in specifically targeting and reducing IL-8 expression, curcumin’s ability to modulate multiple inflammatory pathways provides a complementary and more comprehensive approach. Both strategies show promise, but curcumin’s multi-targeted action offers additional benefits for addressing the complex inflammatory environment in OA.

Figures 8 and 9 represent the qRT-PCR and ELISA results for the primary chondrocytes. The qRT-PCR results for IL-6 and IL-8, represented in Figures 8 (a,c,e) and 9 (a,c,e), respectively, indicate that the mRNA levels in the uninduced cells (NC) were almost negligible in the case of primary chondrocytes (< 0.02 fold change); but they were higher in the C28/I2 cell line (> 0.2 fold change). The higher levels of expression in C28/I2 cells than in the primary cells might be due to the high proliferation rate of the immortalized cell line. In this case, the elevated cytokine mRNA levels observed might also be due to the inherent properties of the immortalized cell line, such as enhanced metabolic activity and stress responses, which can sensitize these cells to inflammatory cues. Immortalized cell lines often show increased baseline cytokine expression as part of their altered physiology, which includes an amplified response to inflammatory signals, but not directly due to cell proliferation alone [58].

**Figure 8.**
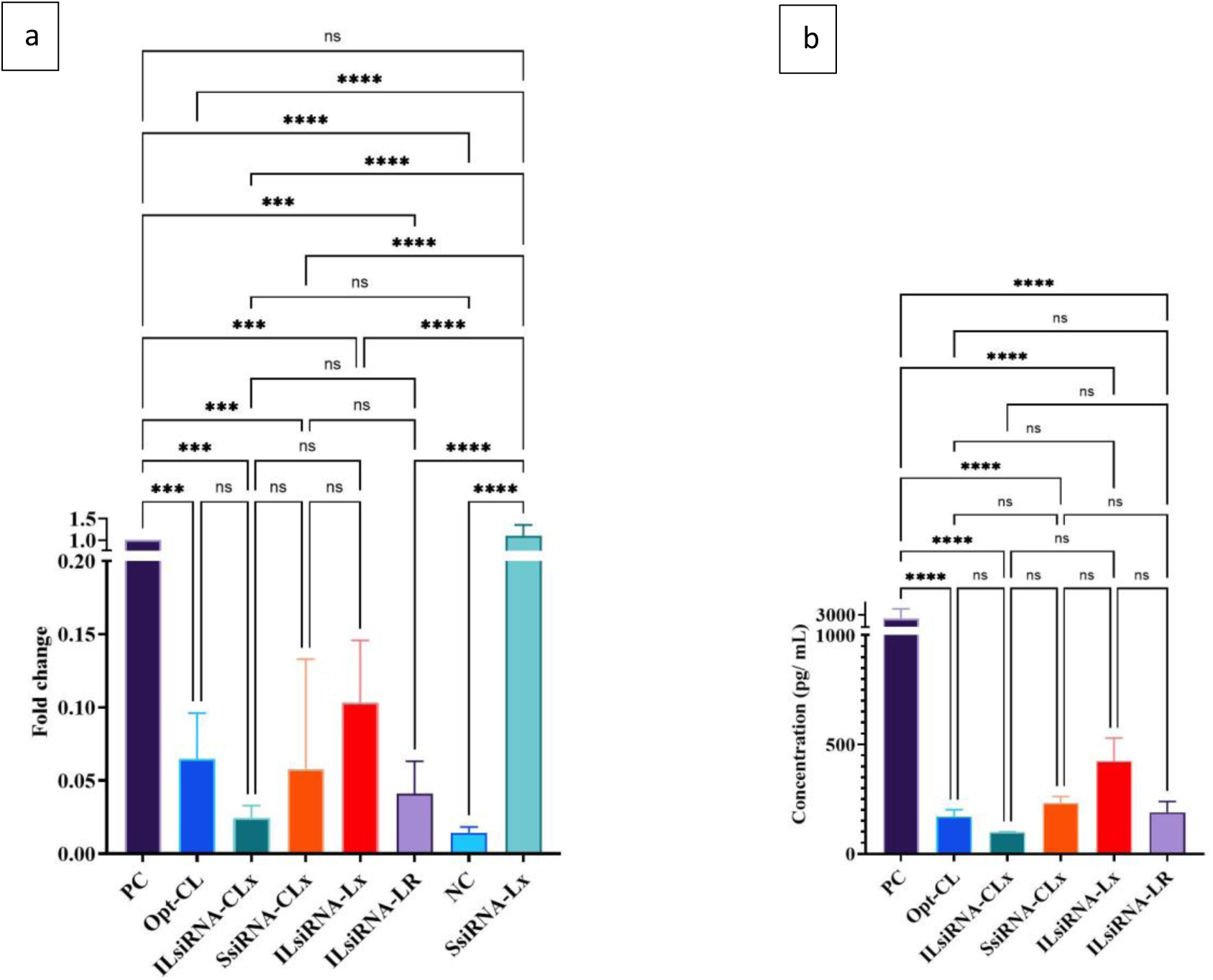
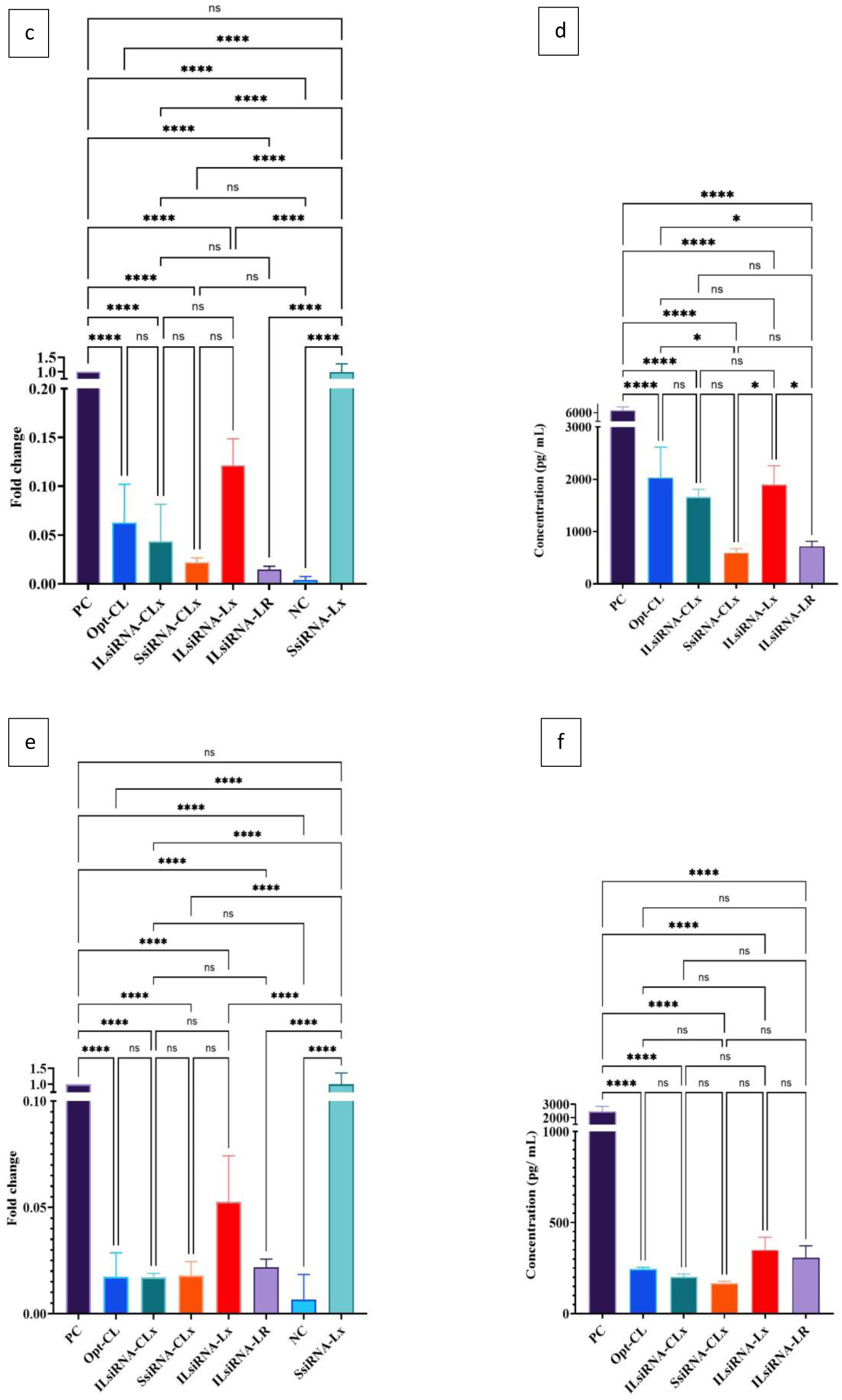
IL-6 mRNA levels were expressed through qRT-PCR (left), and protein levels were quantified using ELISA (right). a&b – primary chondrocytes C28-CA-22-004, c&d – primary chondrocytes C28-CA-22-009, e&f – primary chondrocytes C28-CA-23-004. Positive control (PC), negative control (NC), curcumin-loaded cationic liposomes (Opt-CL), IL-6 and IL-8 siRNA + Curcumin co-loaded lipoplexes (ILsiRNA-CLx), Scramble siRNA + Curcumin co-loaded lipoplexes (SsiRNA-CLx), IL-6 and IL-8 siRNA lipoplexes (ILsiRNA-Lx), Scramble siRNA lipoplexes (SsiRNA-Lx), and IL-6 and IL-8 siRNA with Lipofectamine RNAimax formulation as a standard for transfection and knockdown (ILsiRNA-LR). Results were expressed as mean ± SD of three independent measurements. A one-way ANOVA test with multiple comparisons was performed to analyze the differences between the effects of the treatments (ns, P > 0.05; *, P < 0.05; **, P < 0.01; ***, P < 0.001; ****, P < 0.0001). The graphs were generated using GraphPad Prism 10.2.2.

**Figure 9.**
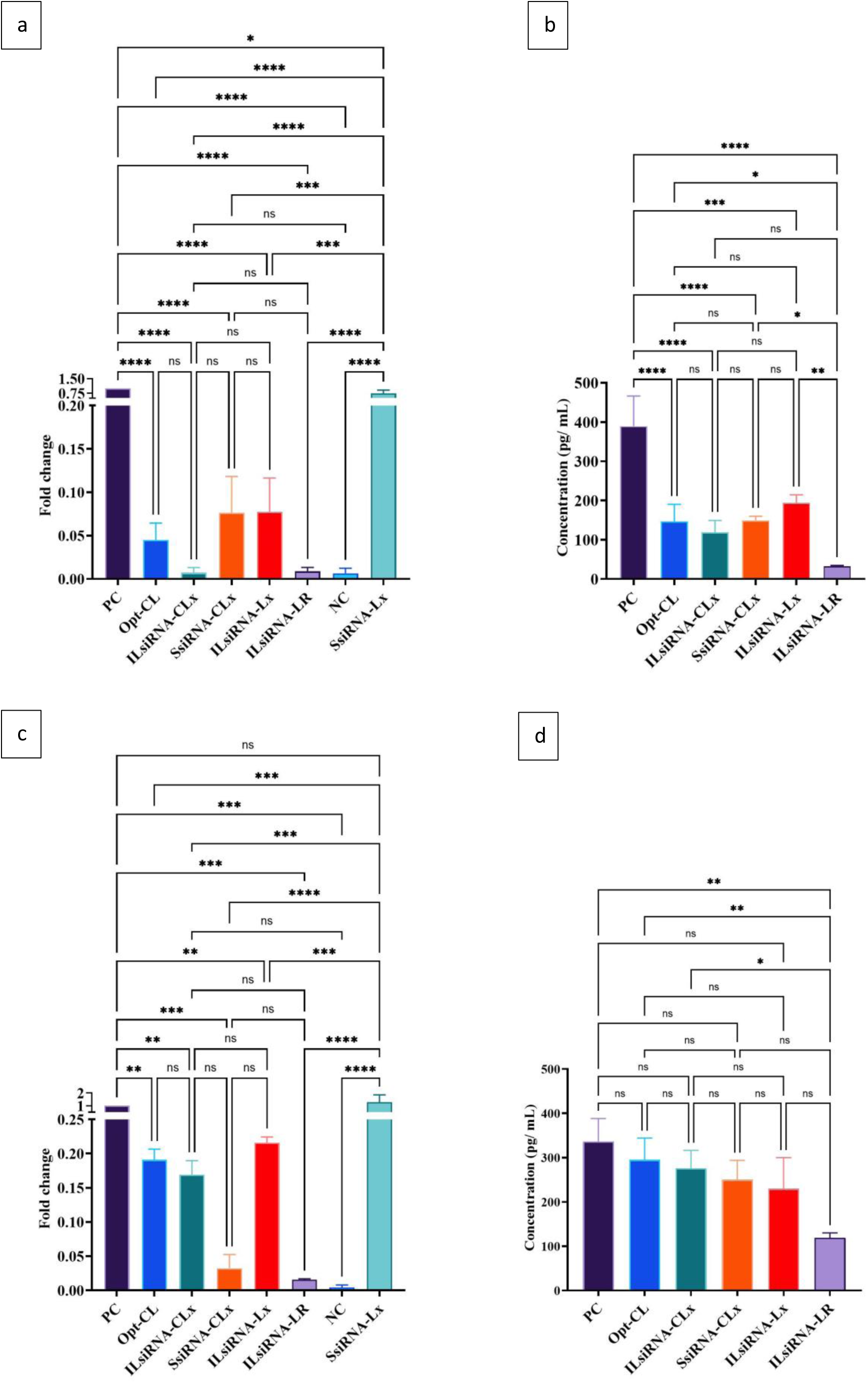
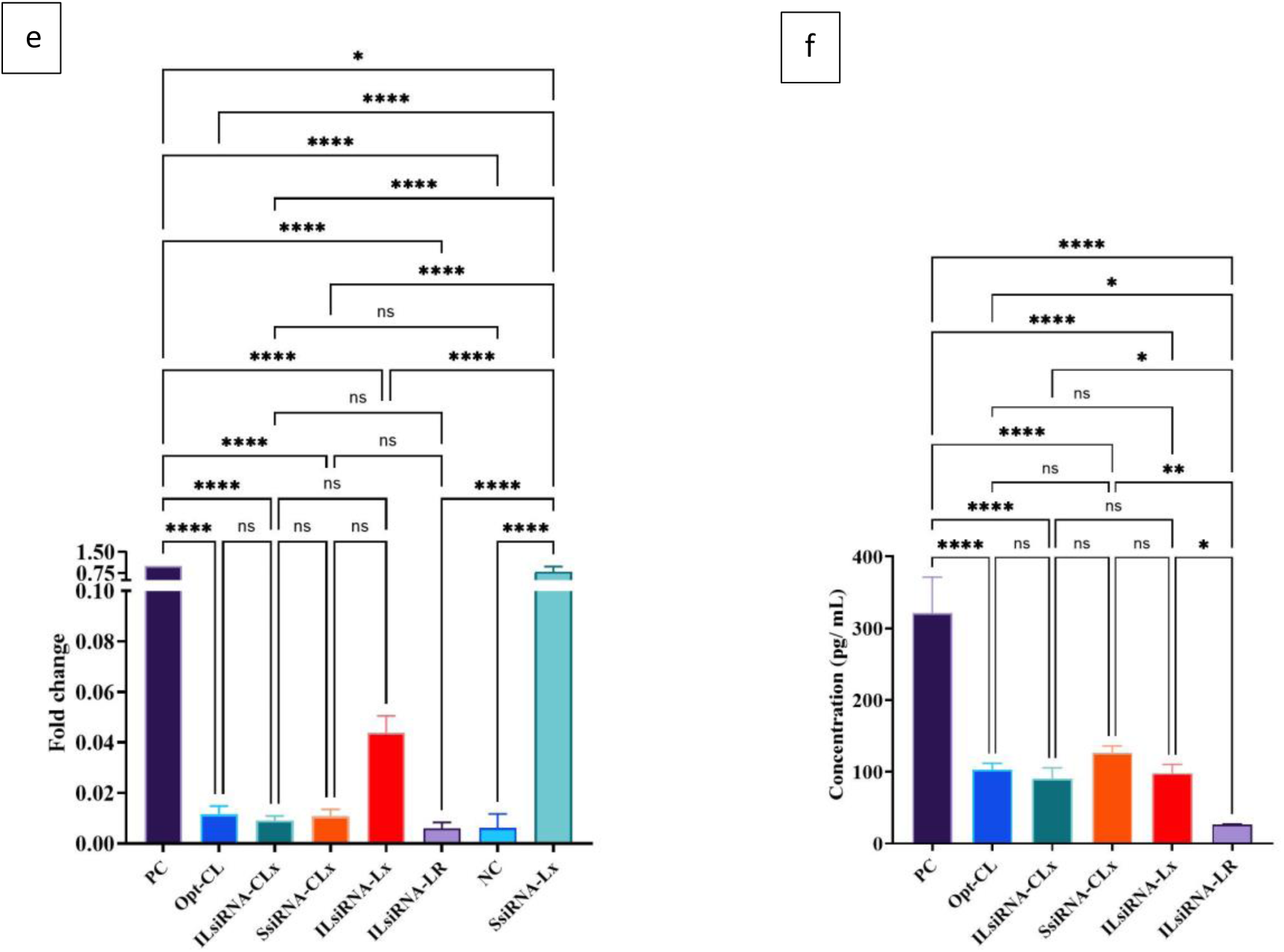
IL-8 mRNA levels were expressed through qRT-PCR (left), and protein levels were quantified using ELISA (right). a&b – primary chondrocytes C28-CA-22-004, c&d – primary chondrocytes C28-CA-22-009, e&f – primary chondrocytes C28-CA-23-004. Positive control (PC), negative control (NC), curcumin-loaded cationic liposomes (Opt-CL), IL-6 and IL-8 siRNA + Curcumin co-loaded lipoplexes (ILsiRNA-CLx), Scramble siRNA + Curcumin co-loaded lipoplexes (SsiRNA-CLx), IL-6 and IL-8 siRNA lipoplexes (ILsiRNA-Lx), Scramble siRNA lipoplexes (SsiRNA-Lx), and IL-6 and IL-8 siRNA with Lipofectamine RNAimax formulation as a standard for transfection and knockdown (ILsiRNA-LR). Results were expressed as mean ± SD of three independent measurements. A one-way ANOVA test with multiple comparisons was performed to analyze the differences between the effects of the treatments (ns, P > 0.05; *, P < 0.05; **, P < 0.01; ***, P < 0.001; ****, P < 0.0001). The graphs were generated using GraphPad Prism 10.2.2.

**Figure 10.**
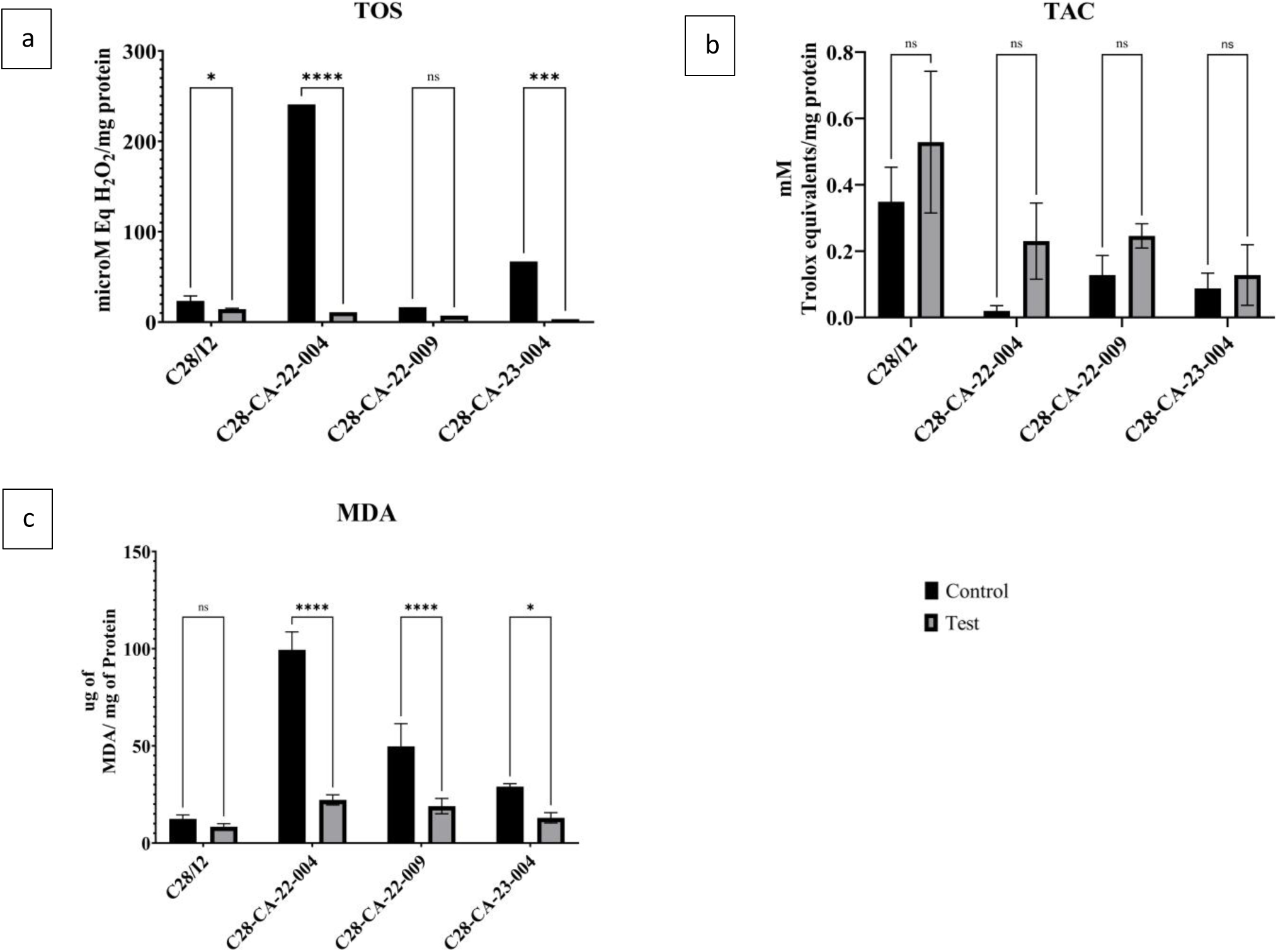
Effects of the administration of Opt-CL on the oxidative stress on C28/I2 cells and primary chondrocytes. Control - H_2_O_2_ induced cells, Test-H_2_O_2_ induced, and liposomes treated cells. A) Total oxidant status (TOS) levels expressed as micromolar equivalents H_2_O_2_/mg protein; B) Total antioxidant capacity (TAC) expressed as mM Trolox equivalents/mg protein; C) Malondialdehyde (MDA) levels (µg) expressed per mg protein. Results were expressed as mean ± SD three independent measurements. A one-way ANOVA test was performed to analyze the differences between the effects of the treatments (ns, P > 0.05; *, P < 0.05; **, P < 0.01; ***, P < 0.001; ****, P < 0.0001). The graphs were generated using GraphPad Prism 10.2.2.

Lipoplexes with only Scramble siRNA (SsiRNA-Lx) had no effect, as the mRNA expression levels were similar (fold change 1) or much higher (fold change greater than 1) than those of the positive control. It appears that almost all the tested liposomal formulations containing either curcumin or siRNA or their combination were significantly effective compared to the induced and untreated cells (positive control). The figures denote that the lipoplexes with siRNA were effective in downregulating the expression of inflammatory cytokines. Moreover, the liposomes containing curcumin either alone (Opt-CL) or combined with siRNA (ILsiRNA-CLx, SsiRNA-CLx) reduced cytokine mRNA expression more effectively than IL-6 and IL-8 siRNA lipoplexes by at least 80%. This influence of curcumin might also give a practical synergistic application in pre-clinical models. The gene silencing activity of the co-loaded lipoplexes (ILsiRNA-CLx) was almost comparable to that of the Lipofectamine standard in primary chondrocytes, and it appears that there is no significant difference between the formulations. Our results indicate that curcumin and siRNA co-loaded lipoplexes specifically and effectively downregulate cytokine gene expression in chondrocytes. The lipoplexes with only siRNA but no curcumin could also effectively reduce the mRNA levels except for the C28-CA-22-009 primary cells.

For the ELISA analysis, the cells treated with Scramble siRNA lipoplexes (SsiRNA-Lx) were not assessed for protein levels, as their mRNA expression in chondrocytes was found to be insignificant. Similarly, the negative control (NC) was not included for simplicity in the ELISA layout design. The ELISA results for other formulations showed some differences compared to the qRT-PCR data. While the ELISA results for IL-6 and IL-8 generally aligned with the mRNA levels obtained from qRT-PCR, the protein levels were higher in magnitude, likely reflecting post-transcriptional and post-translational modifications that mRNA measurements do not capture. This discrepancy is expected, as protein synthesis is influenced by additional regulatory factors beyond transcription [59,60].

In particular, when comparing the positive control group with the test groups, IL-6 and IL-8 levels were notably higher in the C28/I2 cell line compared to primary cells. While the high proliferation rate of the immortalized cell line may partially explain this difference, other factors, such as altered signaling pathways or variations in protein synthesis rates between the two cell types, could have contributed [61]. Additionally, cytokine stability may vary between primary and immortalized cells, further influencing the observed protein levels [62]. Notably, the siRNA-loaded lipoplexes, with or without curcumin (ILsiRNA-CLx, ILsiRNA-Lx), more effectively decreased the protein expression for both IL-6 and IL-8 compared to other treatments, as shown in Figures 8 and 9.

For IL-6, the ILsiRNA-CLx markedly affected the protein expression levels compared to ILsiRNA-Lx in primary chondrocytes. The extent of reduction in the protein expression levels was almost similar to the treatment with Lipofectamine, known for its high transfection efficiency. However, this effect was only considerable in primary chondrocytes, and there was no significance for the C28/I2 cells. The protein levels of IL-8 in both the C28/I2 cell line and patient-derived C28-CA-22-009 cells showed minimal change after treatment, with levels remaining similar to the positive control. While high cell division rates in the C28/I2 cells might partly explain this, patient-derived cells like C28-CA-22-009 likely exhibit a more complex interaction of factors. These cells, being closer to a physiological state, could have baseline alterations in inflammatory signaling pathways due to the disease state of the donor. This could result in a less responsive behavior to treatments targeting IL-8, as the inflammatory environment in diseased cells is often harder to modulate. Additionally, in both cell types, factors such as persistent NF-κB activation and post-translational regulatory mechanisms might maintain elevated IL-8 levels, reducing the impact of the treatments. For patient-derived cells, cytokine stability or altered degradation pathways might further contribute to sustained IL-8 protein levels, similar to what is observed in immortalized cell lines.

In the other two patient cells, even though the formulation with Lipofectamine produced the best results regarding IL-8 protein levels, the tested formulations also could significantly reduce the protein levels compared to the positive control, not only for IL-8 (>50%), but also for IL-6 (>80%). Drawing from these results, the co-loaded lipoplexes with curcumin and the IL6 and IL8 siRNA effectively knocked down the inflammatory cytokine genes and significantly reduced protein levels.

These observations confirmed the previous findings of Zhang et al. and Nicoliche et al., which stated that curcumin slowed the progression of OA and exerted a protective effect on the cartilage [63,64]. Gorabi et al., in their meta-analysis of randomized controlled trials on the effect of curcumin on pro-inflammatory cytokines, showed that curcumin significantly reduced the pro-inflammatory cytokines IL-1 and TNF-α and not necessarily IL-6 and IL-8 [65]. So, the delivery system developed in this study allowed a more complex inhibitory activity on pro-inflammatory markers, IL-6 and IL-8 overexpressed in OA. However, the liposomal formulations in this study effectively delivered the curcumin and silenced the genes for the inflammatory cytokines. These co-loaded lipoplexes are an effective carrier for both the oligonucleotides and the drug molecules. When a long-term treatment is required for conditions like OA, potent drugs like curcumin could be delivered for prolonged periods using liposomes. These formulations could be further used to be evaluated in pre-clinical and clinical models for more data correlation, which is finally destined to help the patients suffering from OA. This approach of formulating the co-loaded lipoplexes is so versatile that it could be applied to deliver other hydrophobic drug molecules and small oligonucleotides for various disease models.

### 3.4. Antioxidant effects by curcumin loaded in *Opt-CL* on human chondrocytes

To elucidate the antioxidative potential of our therapeutic approach, we employed an *in vitro* model wherein chondrocytes were subjected to oxidative stress induced by hydrogen peroxide (H_2_O_2_) [7,66–68]. Subsequent assessment of cellular oxidative status by comprehensive analysis of key biomarkers, including TAC, TOS, and MDA levels, provided insights into the efficacy of curcumin in mitigating oxidative damage and restoring redox balance [33]. These assessments were performed immediately after oxidative stress induction and protein concentration was used to normalize the quantification of the measurements across different cell lysates with the protein content. The cells that H_2_O_2_ induced were termed control, and the cells treated with opotimum curcumin-loaded liposomes were termed test.

Upon induction of oxidative stress with H_2_O_2_, TOS levels increased significantly for control, reflecting heightened oxidative activity (C28/I2 cells – 23.39 ± 5.54 microM Eq H_2_O_2_/mg protein, primary chondrocytes – 16.52 to 240.94 microM Eq H_2_O_2_/mg protein). TOS levels in chondrocytes treated with Opt-CL (C28/I2 cells – 14.29 ± 0.95 microM Eq H_2_O_2_/mg protein, primary chondrocytes – 3.29 to 10.87 microM Eq H_2_O_2_/mg protein) were significantly lower (P<0.05) than for the control cells, as shown in Figure 10 (a). These results from the amounts of H_2_O_2_ detected in the control and treated cells convey that the Opt-CL reduced cell oxidation to a greater extent due to their antioxidant capacity, as established in the literature [69–72].

Unlike TOS, when the cells were oxidative stress-induced using H_2_O_2_, the TAC test revealed lower levels of the Trolox equivalents (C28/I2 cells – 0.35 ± 0.10 mM Trolox equivalent/mg of protein, primary chondrocytes – 0.02 to 0.13 mM Trolox equivalent/mg of protein) for control. The TAC levels in chondrocytes treated with Opt-CL showed improvement (C28/I2 cells: 0.53 ± 0.21 mM Trolox equivalent/mg of protein; primary chondrocytes: 0.13 to 0.25 mM Trolox equivalent/mg of protein), as shown in Figure 10 (b). However, the increase in antioxidant levels was not statistically significant (P > 0.05). This signifies that a higher amount of Trolox underwent oxidation for the control cells than those treated with Opt-CL. Seemingly, the antioxidant capacity of the test cells treated with Opt-CLwas superior to that of the control, which is in accordance with the reported literature [69,70,73]. It can also be observed that the TAC for the C28/I2 cell line was significantly higher compared to the primary chondrocytes, possibly due to a higher proliferation rate of the former. Primary chondrocytes are exposed to relatively low oxygen concentrations within cartilage tissue. When cultured, they are exposed to higher oxygen levels, which can increase their susceptibility to oxidative damage. Compared to primary chondrocytes, the C28/I2 cells are immortalized by the introduction of a viral oncogene (SV40 large T-antigen) that alters signaling pathways controlling cellular stress and stress responses, providing them with greater tolerance to oxidative damage [74].

Similar to the TOS, upon induction of oxidative stress by H_2_O_2_, the control cells contained high levels of MDA (C28/I2 cells – 12.40 ± 2.05 µg of MDA/mg of protein, primary chondrocytes – 29.04 to 99.41 µg of MDA/mg of protein). Upon the treatment of oxidative stress-induced cells with the Opt-CL, it appears that in the C28/I2 cell line, the levels of MDA did not vary much (P > 0.05); however, the primary chondrocytes from all three patients showed significant differences in MDA levels compared to control (P < 0.05), as shown in Figure 10 (c). This might suggest that Opt-CL reduced the lipid peroxidation levels [75–77].

## 4. Conclusion

We successfully applied a QbD approach to optimize co-loaded lipoplexes containing curcumin and siRNAs against inflammatory cytokines involved in OA pathology. The QTPP was defined, guiding the identification of CQAs, CMAs, and CPPs. The CQAs for the liposomal formulation included PS, PDI, ZP, EE, cell viability, and transfection. Risk assessment identified lipid composition, cationic lipid concentration, PEGylated phospholipid, and cholesterol content as critical variables for the safety and efficacy of the liposomal formulations. Thus, using DoE, the liposomal formulation was designed and optimized with regard to its stability, curcumin release, and cell uptake of siRNA to ensure efficient and simultaneous suppression of oxidative stress and inflammation associated with OA. Thus, a liposomal formulation with a lipid composition of 8 mM DOPE, 2 mM MPEG-DSPE, 5 mM DOTAP, and 5 mM cholesterol was stable for 6 months at 4°C and provided a prolonged curcumin release for 72 hours *in vitro*.

Moreover, the formulation achieved comparable transfection efficiency to Lipofectamine RNAiMAX, effectively reducing IL-6 and IL-8 mRNA and protein levels in the human articular chondrocytes. Furthermore, *in vitro* studies on human chondrocytes (C28/I2 cell line and articular chondrocytes) demonstrated the potential of curcumin- and siRNA-loaded lipoplexes to reduce oxidative stress due to the antioxidant capacity of curcumin as well as inflammation-driven by IL-6 and IL-8 due to the siRNAs internalized efficiently in these cells. These findings highlighted the potential of curcumin- and siRNA-loaded lipoplexes for intra-articular administration, offering prolonged therapeutic release and enhanced patient comfort.

Given these promising results, further exploration of these co-loaded lipoplexes as combination or adjuvant therapies for OA is warranted. Their potential for use alongside biomaterials like hyaluronic acid conjugates or as a complement to existing OA therapies could provide a targeted and efficient means of reducing inflammation and oxidative stress, improving outcomes for OA patients.

## CRediT author statement

**Saketh Reddy Ranamalla –** Conceptualization, Methodology, Validation, Formal analysis, Investigation, Data Curation, Writing - Original Draft, Visualization, **Alina Porfire –** Methodology, Validation, Writing - Review & Editing, **Emilia Licarete –** Methodology, Validation, Writing - Review & Editing, **Lucia Tefas –** Writing - Review & Editing, **Rohith Pavan Parvathaneni –** Methodology, Validation, Formal analysis, Investigation, Data Curation**, Alina Sesarman** – Writing - Review & Editing, Resources, **Monica Focsan –** Methodology, Investigation, Data Curation, **Lucian Barbu Tudoran** – Methodology, Investigation, Data Curation, **Oommen. P. Varghese –** Writing - Review & Editing, Resources, Supervision, **Ioan Tomuta –** Writing - Review & Editing, Resources, Supervision, Project administration, Funding acquisition, **Manuela Banciu –** Writing - Review & Editing, Resources, Supervision.

## Acknowledgments

The C28/I2 cells were a kind gift from Prof Mary Goldring, Professor of Cell and Developmental Biology at Weill College Graduate School of Medical Sciences, New York, United States. The authors would also like to acknowledge Prof. Laura Creemers and Ms. Katrin Agnes Münzebrock, UMC Utrecht, The Netherlands, for their support in providing the primary chondrocytes for the *in vitro* cell studies. The authors would also like to thank Adorján Cristea and Bogdan Dume Razvan of the Faculty of Biology and Geology, “Babes-Bolyai” University, for supporting the *in vitro* cell experiments.

