## Supplemental file for "A QbD strategy to develop curcumin and siRNA co-loaded lipoplexes to combat osteoarthritis-related inflammation and oxidative stress"

* Corresponding authors

### Synthesis of luciferase siRNA

Luciferase siRNA was initially used as a prototype oligonucleotide material to optimize the cationic liposomes. It was used to silence the expression of the luciferase gene in the luciferase-expressing C28/I2 cell line. Luciferase siRNA Sense (5' rCrUrUrArCrGrCrUrGrArGrUrArCrUrUrCrGrAdTdT 3') and Antisense (5' rUrCrGrArArGrUrArCrUrCrArGrCrGrUrArArGdTdT 3') were synthesized using solid phase oligonucleotide synthesis by H8 K&A synthesizer with standard 2'-O-TBDMS protected monomers through phosphoramidite chemistry. Cleavage from solid support and deprotection of cyanoethyl and nucleotide protecting groups was done by adding 1 mL (1:1 v/v) of ammonium hydroxide and methylamine (AMA) solution to solid beads at 65°C. 2'-O-TBDMS was deprotected by treating with triethylamine trihydrofluoride, and the reaction was quenched with isopropoxytrimethylsilane. The crude oligonucleotide was precipitated by ether, purified by 20% denaturing PAGE (7M Urea), and recovered with Tris-EDTA-NaCl (TEN) buffer. RNA samples were desalted using Waters (WAT020515) Sep-Pak column. The pure RNA pellet was dissolved in water, and the concentration was measured at 260 nm using a Cary 3500 Multicell UV-Vis Spectrophotometer. A duplex was formed by adding an equimolar ratio of sense and antisense strands through heating at 90°C for 10 min and gradually cooling to room temperature. MALDI analysis was performed at BianoGMP GmbH. The MALDI analysis is shown in Figure S.1.

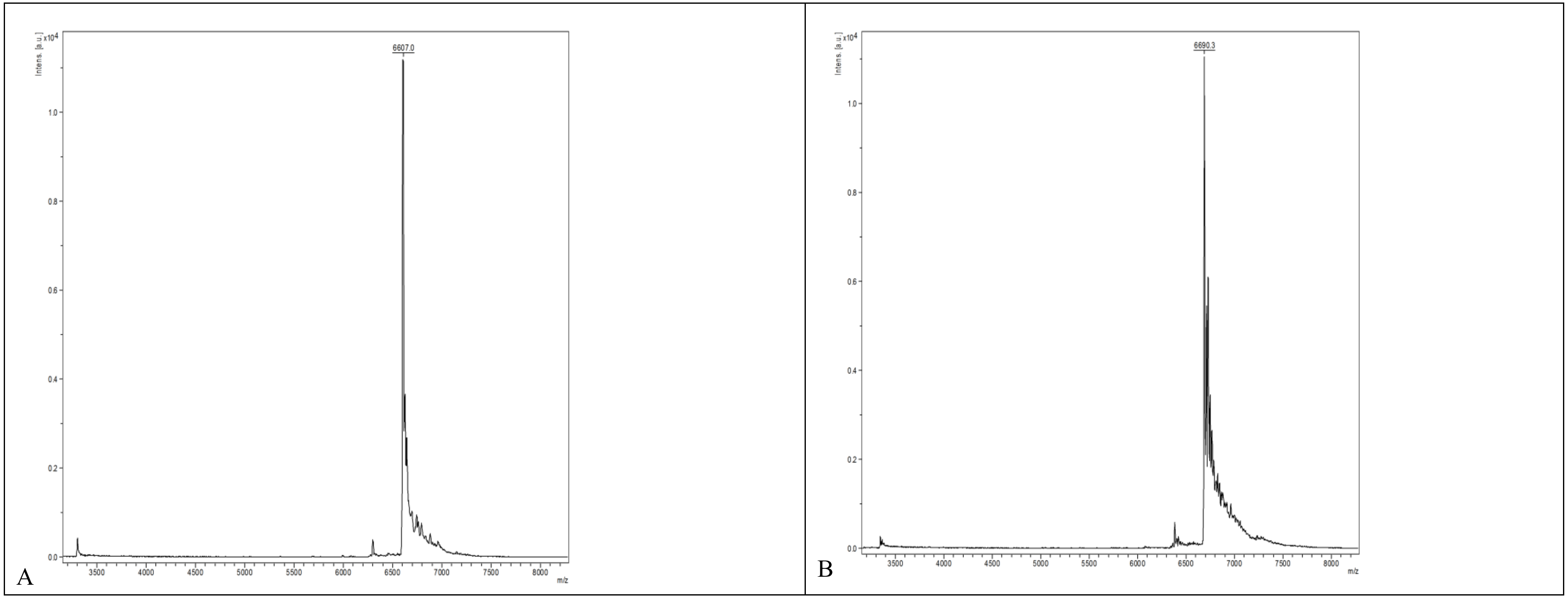

**Figure S.1.** *MALDI TOF analysis of Sense strand (left - Theoretical Mass: 6607 g/mol) and Antisense strand (right - Theoretical Mass: 6693 g/mol) of Luciferase siRNA*

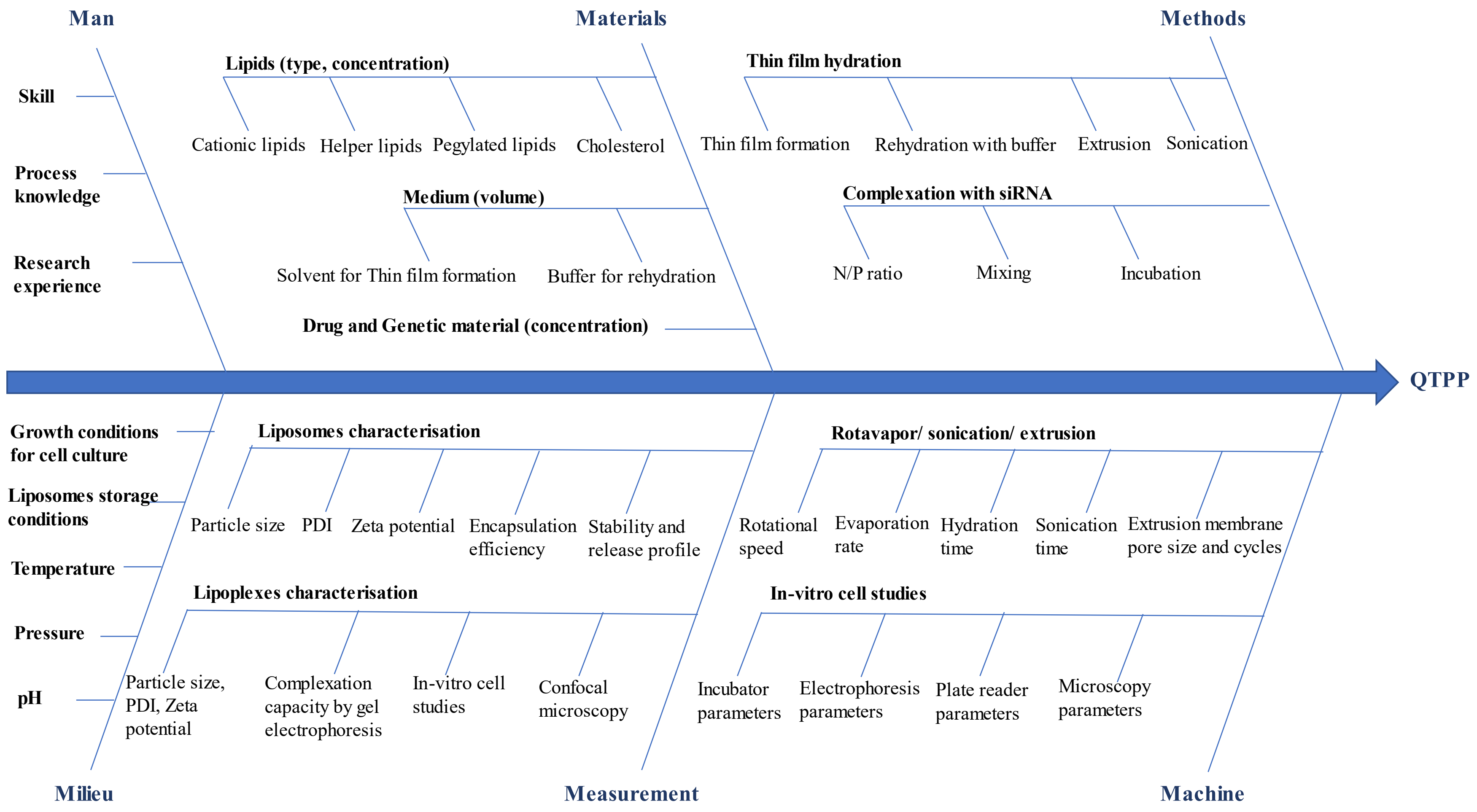

**Figure S.2.** *Ishikawa diagram for the preparation of co-loaded lipoplexes for intra-articular delivery*

**Table S.1.** FMEA analysis for the co-loaded lipoplexes

| Parameter | Failure Mode | Effects | Cause | O | S | D | RPN | Mitigation Strategy |
|---|---|---|---|---|---|---|---|---|
| **Formulation parameters** | | | | | | | | |
| Lipid Composition | Incorrect cationic lipid: helper lipid ratio | Poor encapsulation efficiency, unstable liposomes | Miscalculation, improper mixing | 3 | 9 | 5 | 135 | Standardize mixing protocols, precise analytical balance usage, and validation of lipid ratios. |
| **Lipid Composition** | **Inadequate cationic lipid content** | **Reduced gene delivery efficiency** | **Misinterpretation of formulation requirements** | **3** | **9** | **6** | **162** | **Optimize lipid ratio with experimental studies and review formulation protocol regularly.** |
| **Curcumin Loading** | **Low encapsulation efficiency** | **Reduced therapeutic efficacy** | **Suboptimal solvent, lipid solubility, mixing technique** | **4** | **8** | **5** | **160** | **Optimize solvent system, adjust mixing conditions, and conduct optimization trials for curcumin loading.** |
| Particle Size | Too large or too small particles | Reduced cell uptake, inefficient gene delivery | Inadequate sonication/extrusion parameters | 3 | 9 | 4 | 108 | Standardize sonication/extrusion conditions and monitor particle size distribution regularly. |
| **Surface Charge** | **Insufficient positive charge** | **Decreased interaction with cell membranes** | **Inadequate cationic lipid, lipid oxidation** | **4** | **9** | **5** | **180** | **Check lipid quality, adjust cationic lipid ratio, use antioxidants** |
| Curcumin Stability | Degradation during formulation | Loss of efficacy, potential toxicity | Light, oxygen exposure, inappropriate pH | 3 | 8 | 6 | 144 | Light and oxygen protection, pH optimization, stabilizer addition |
| Gene Encapsulation | Low encapsulation efficiency | Ineffective gene silencing, reduced therapeutic effect | Improper complexation conditions, nucleic acid degradation | 3 | 9 | 5 | 135 | Optimize nucleic acid to lipid ratio, gentle mixing, and nucleic acid protection during formulation. |

| Parameter | Failure Mode | Effects | Cause | O | S | D | RPN | Mitigation Strategy |
|---|---|---|---|---|---|---|---|---|
| **Process parameters - Liposome preparation** | | | | | | | | |
| Thin Film Formation | Incomplete dissolution of lipids | Heterogeneous liposome populations, varying sizes | Inadequate solvent choice or volume, improper mixing | 3 | 7 | 5 | 105 | Optimize solvent selection to ensure complete mixing |
| Hydration of Thin Film | Inefficient hydration | Low encapsulation efficiency | Inadequate hydration time, temperature, or medium | 3 | 7 | 5 | 105 | Optimize hydration parameters (time, temperature, medium) |
| Sonication | Excessive or insufficient sonication | Liposome destabilization or inadequate size reduction | Incorrect sonication intensity or duration | 4 | 7 | 5 | 140 | Standardize sonication parameters, calibrate equipment, monitor process |
| Extrusion | Membrane clogging or inconsistent pressure | Variable liposome size distribution | Inappropriate membrane pore size, inadequate filtration, improper operation | 3 | 9 | 4 | 108 | Select appropriate membrane filters and follow operational guidelines. |
| Post-Extrusion Particle Size Analysis | Inaccurate size measurement | Potential efficacy loss | Malfunctioning or improperly calibrated equipment | 2 | 7 | 5 | 70 | Regular calibration and maintenance of equipment |
| **Stability** | **Oxidative degradation of lipids** | **Compromised liposome integrity, premature release** | **Exposure to oxygen, improper storage conditions** | **4** | **9** | **6** | **216** | **Use proper packaging/ store under nitrogen/ appropriate storage temperature.** |
| Stability | Hydrolysis of lipids | Altered membrane fluidity, destabilized liposomes | Moisture, high temperature | 3 | 8 | 5 | 120 | Control storage conditions (temperature, humidity) |

| Parameter | Failure Mode | Effects | Cause | O | S | D | RPN | Mitigation Strategy |
|---|---|---|---|---|---|---|---|---|
| Stability | Phase separation | Heterogeneous drug distribution, reduced efficacy | Incompatibility of lipid components, temperature fluctuations | 2 | 7 | 5 | 70 | Optimize lipid composition, ensure uniform mixing, and stabilize storage conditions. |
| Release | Rapid curcumin release | Reduced therapeutic window, increased side effects | High membrane fluidity, inappropriate pore size | 4 | 9 | 4 | 144 | Adjust lipid composition for controlled release |
| Release | Incomplete curcumin release | Decreased efficacy, wasted drug | Too rigid membrane structure, improper encapsulation technique | 3 | 9 | 5 | 135 | Optimize lipid-to-drug ratio and optimize hydration and sonication conditions. |
| **Release** | **curcumin degradation** | **Loss of efficacy, potential toxic degradation products** | **Light exposure, temperature, pH** | **4** | **8** | **6** | **192** | **Protect from light, control pH and temperature during storage and handling** |
| **Process parameters - Lipoplex preparation** | | | | | | | | |
| **Lipid- RNA Ratio** | **Incorrect lipid-to-nucleic acid ratio** | **Inefficient complex formation, reduced transfection efficiency** | **Miscalculation, inaccurate measurement** | **4** | **9** | **5** | **180** | **Standardize measurement techniques, validate lipid-to-nucleic acid ratio, and conduct preliminary optimization studies.** |
| Lipid Quality | Poor-quality lipid reagents | Decreased transfection efficiency, cytotoxicity | Contaminants, degradation of lipid components | 3 | 9 | 6 | 162 | Source high-quality reagents and store lipids properly. |

| Parameter | Failure Mode | Effects | Cause | O | S | D | RPN | Mitigation Strategy |
|---|---|---|---|---|---|---|---|---|
| Buffer Composition | Inappropriate buffer conditions | Altered lipoplex formation, reduced stability | Incorrect pH, ionic strength, or buffer components | 3 | 8 | 5 | 120 | Optimize buffer conditions based on literature or preliminary experiments to validate buffer composition. |
| Mixing Parameters | Inadequate mixing parameters | Inefficient complexation, non-uniform particle size distribution | Insufficient mixing time, speed, or method | 4 | 8 | 4 | 128 | Standardize mixing parameters, use validated protocols, and optimize mixing conditions. |
| Nucleic Acid Quality | Degraded or impure nucleic acids | Reduced transfection efficiency, potential toxicity | Degradation during handling or purification, contamination | 3 | 9 | 5 | 135 | Source high-quality nucleic acids, handle and store nucleic acids properly |
| **Cationic Lipid Toxicity** | **Cytotoxicity of cationic lipids** | **Cell death, reduced transfection efficiency** | **Non-specific cell membrane disruption, interaction with serum proteins** | **4** | **9** | **6** | **216** | **Perform cytotoxicity assays, optimize lipid-to-nucleic acid ratio to minimize toxicity, and use lipids with lower toxicity profiles.** |
| **Cell culture study aspects** | | | | | | | | |
| **Cell Viability** | **Cytotoxicity induced by lipoplexes** | **Cell death, compromised study outcomes** | **Toxic components in lipoplex formulation, excessive concentration, or improper handling** | **4** | **9** | **6** | **216** | **Optimize lipoplex formulation for reduced toxicity, control dosage and exposure time, and validate results with multiple assays or replicate experiments.** |
| Cell Viability | Interference with cell viability assays | False positive or negative results, inaccurate assessment of lipoplex effects | Components of lipoplexes interfering with assay reagents, optical interference, or detection limits | 3 | 8 | 5 | 120 | Validate compatibility of lipoplex components with assay reagents, optimize assay conditions for sensitivity and specificity, and include appropriate controls to account for potential interference. |

| Parameter | Failure Mode | Effects | Cause | O | S | D | RPN | Mitigation Strategy |
|---|---|---|---|---|---|---|---|---|
| **Cell Internalization** | **Inefficient cellular uptake of lipoplexes** | **Reduced transfection efficiency, inaccurate assessment of cellular effects** | **Inappropriate particle size, surface charge, or composition, ineffective transfection agents** | **4** | **9** | **6** | **216** | **Optimize lipoplex formulation for enhanced cellular uptake, optimize incubation time and conditions, and validate internalization using microscopy techniques or flow cytometry.** |
| Cell Internalization | Non-specific binding and aggregation of lipoplexes | Altered cellular response, misinterpretation of uptake mechanisms | Aggregation due to improper formulation, non-specific binding to cell surface receptors | 3 | 8 | 5 | 120 | Optimize lipoplex formulation to minimize aggregation, validate internalization using microscopy techniques and consider alternative methods such as confocal microscopy or TEM for detailed visualization and quantification of internalization. |
| **Cell Transfection** | **Inefficient gene delivery and expression** | **Reduced expression of target genes, inaccurate assessment of lipoplex efficacy** | **Suboptimal lipoplex formulation, ineffective transfection agents, low transfection efficiency** | **4** | **9** | **6** | **216** | **Optimize lipoplex formulation for improved transfection efficiency, validate transfection efficiency using reporter genes or qPCR, and utilize positive controls to assess transfection efficiency.** |
| Cell Transfection | Non-specific gene expression | Off-target effects, misinterpretation of specific gene effects | Non-specific binding of lipoplexes to non-target cells, off-target gene expression | 3 | 8 | 5 | 120 | Optimize lipoplex formulation for target specificity and use appropriate controls to validate specificity. |

O-Occurrence, S-Severity, D-Detectability, RPN-Risk Priority Number

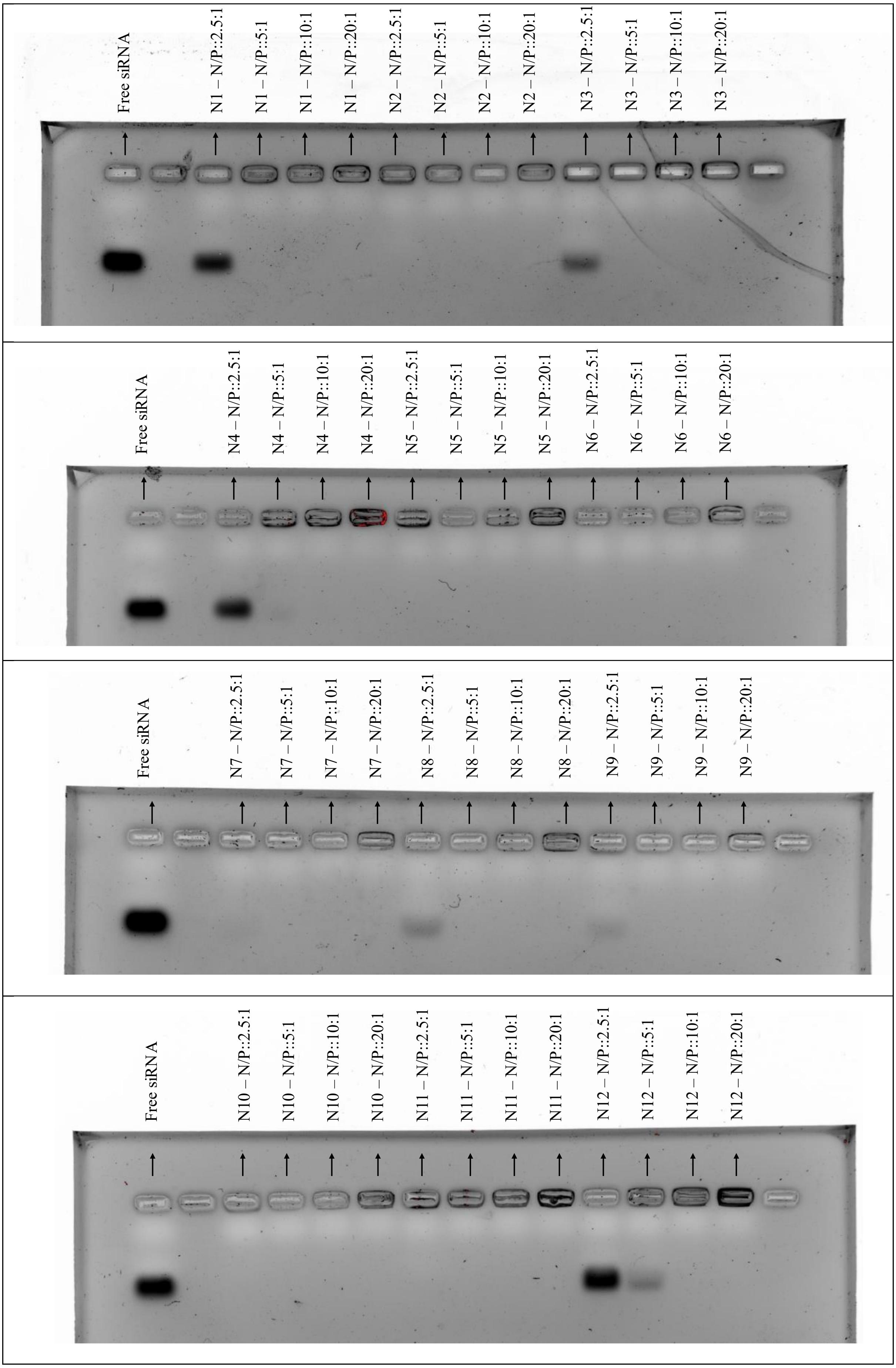
Free siRNA
N1 – N/P::2.5:1
N1 – N/P::5:1
N1 – N/P::10:1
N1 – N/P::20:1
N2 – N/P::2.5:1
N2 – N/P::5:1
N2 – N/P::10:1
N2 – N/P::20:1
N3 – N/P::2.5:1
N3 – N/P::5:1
N3 – N/P::10:1
N3 – N/P::20:1
Free siRNA
N4 – N/P::2.5:1
N4 – N/P::5:1
N4 – N/P::10:1
N4 – N/P::20:1
N5 – N/P::2.5:1
N5 – N/P::5:1
N5 – N/P::10:1
N5 – N/P::20:1
N6 – N/P::2.5:1
N6 – N/P::5:1
N6 – N/P::10:1
N6 – N/P::20:1
Free siRNA
N7 – N/P::2.5:1
N7 – N/P::5:1
N7 – N/P::10:1
N7 – N/P::20:1
N8 – N/P::2.5:1
N8 – N/P::5:1
N8 – N/P::10:1
N8 – N/P::20:1
N9 – N/P::2.5:1
N9 – N/P::5:1
N9 – N/P::10:1
N9 – N/P::20:1
Free siRNA
N10 – N/P::2.5:1
N10 – N/P::5:1
N10 – N/P::10:1
N10 – N/P::20:1
N11 – N/P::2.5:1
N11 – N/P::5:1
N11 – N/P::10:1
N11 – N/P::20:1
N12 – N/P::2.5:1
N12 – N/P::5:1
N12 – N/P::10:1
N12 – N/P::20:1

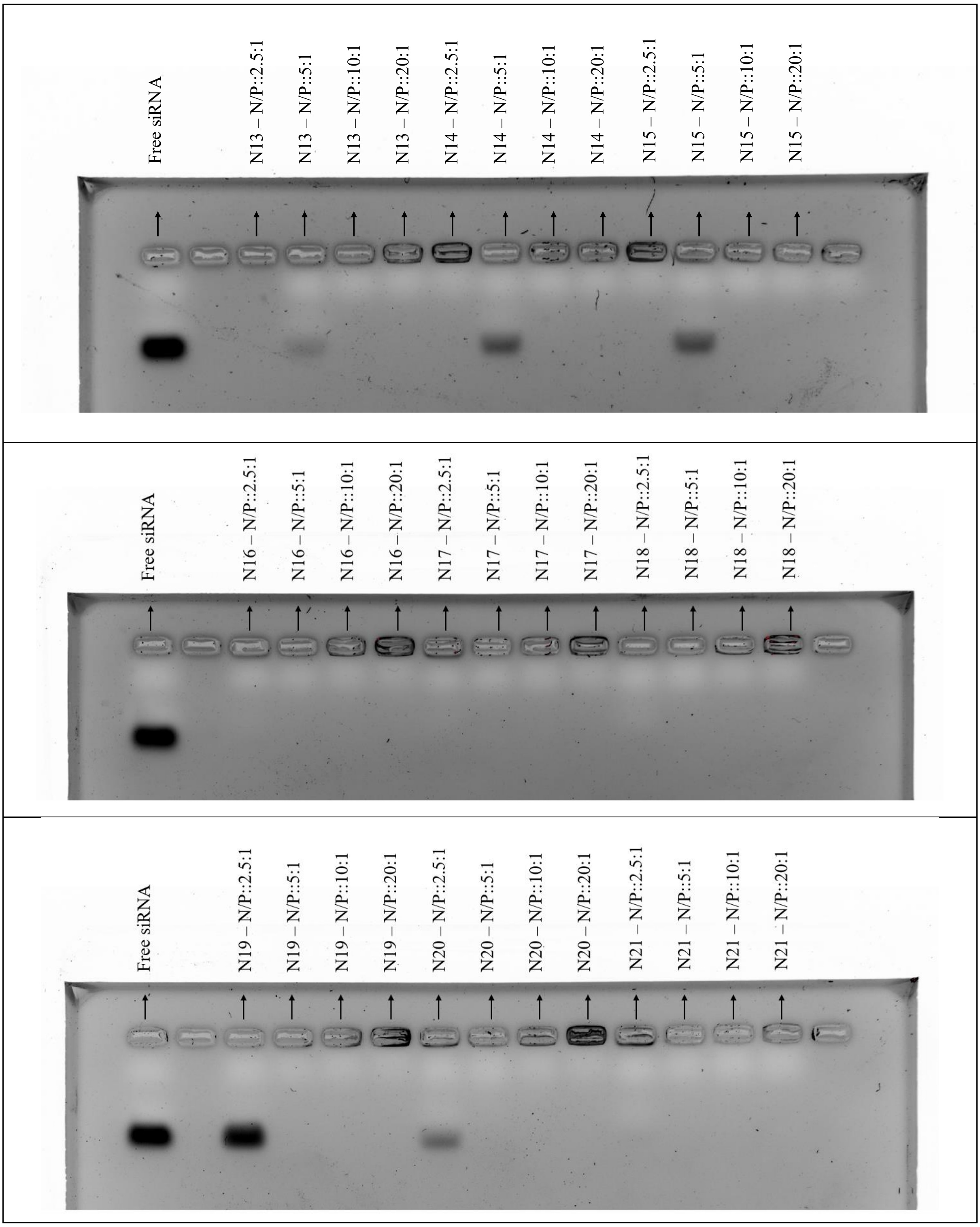

**Figure S.3.** *Complexation capacity measurement by gel electrophoresis and nucleic acid dye staining. N/P refers to the Nitrogen to Phosphate ratio*

| | Cationic liposomes with DOPE | Cationic liposomes with DPPC | Cationic liposomes with DOPC |
|---|---|---|---|
| Visual | 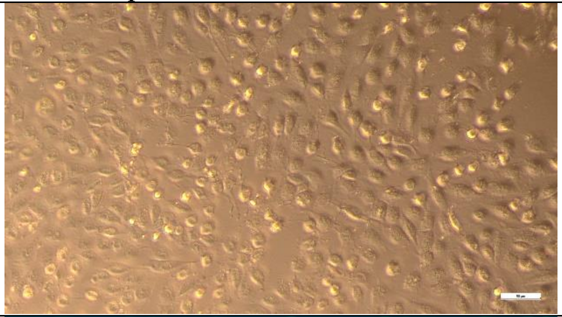 | 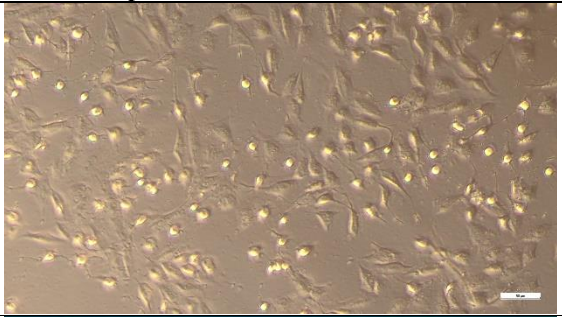 | 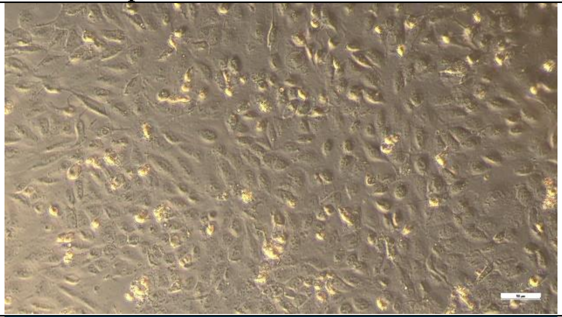 |
| Fluorescence | 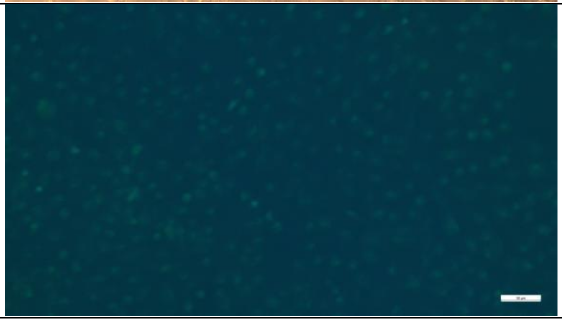 | 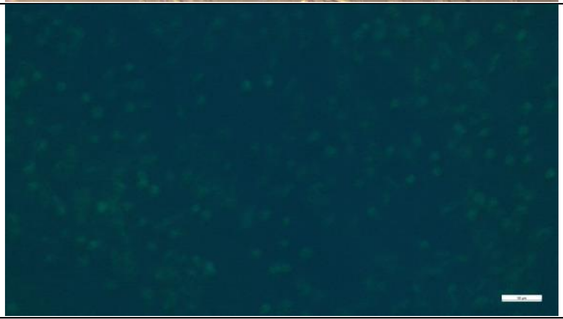 | 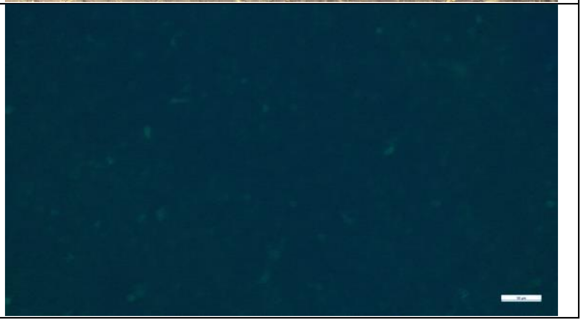 |

**Figure S.4.** *Fluorescence images of C28/I2 cells transfected with curcumin-loaded liposomes*

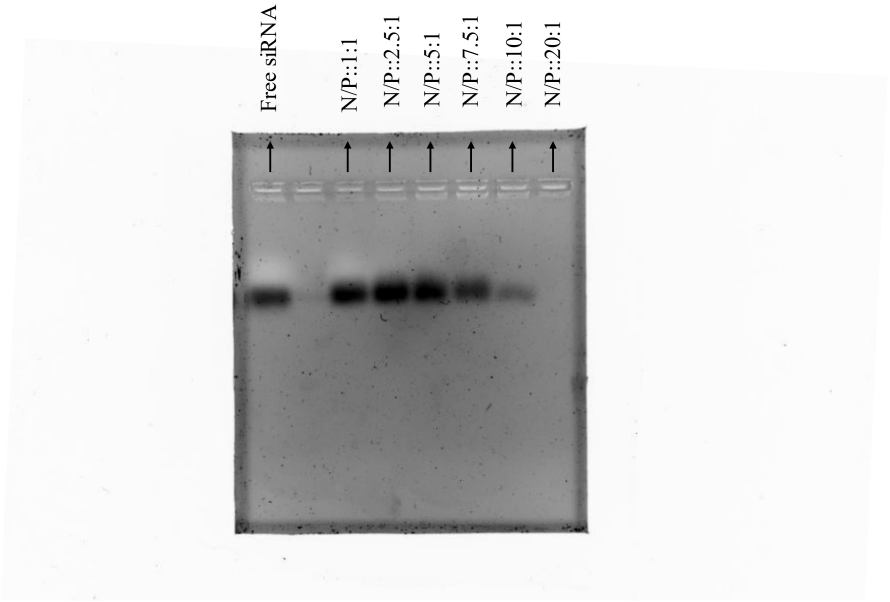

**Figure S.5.** *Complexation capacity for the optimum liposomes' formulation*

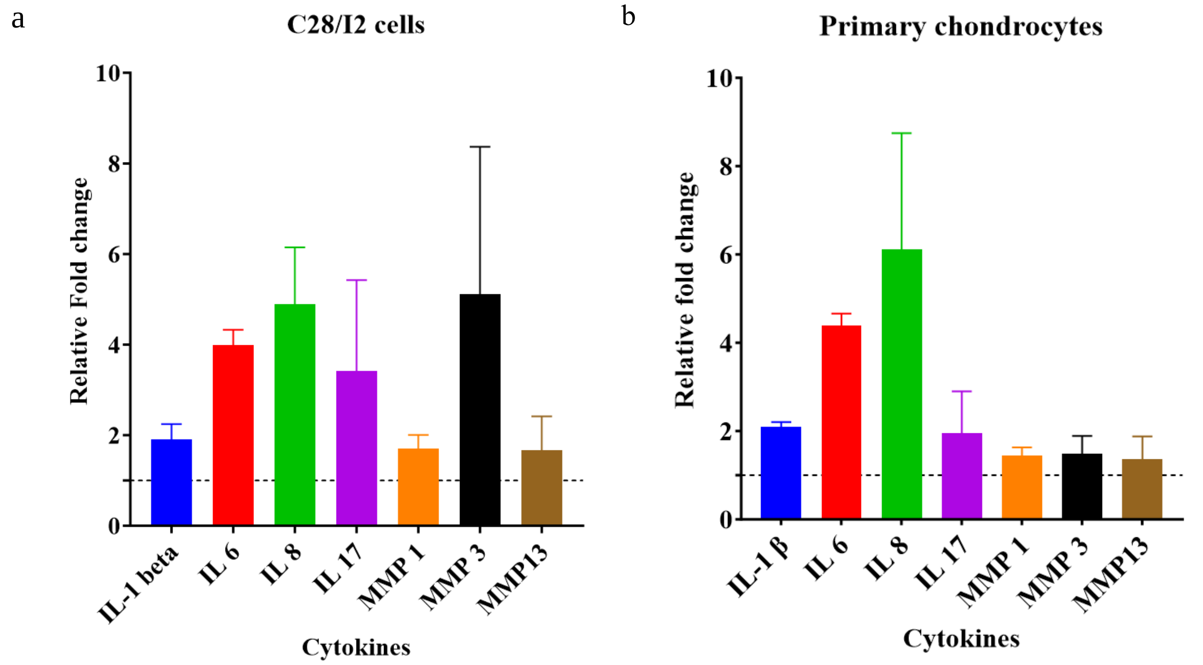

**Figure S.6.** *Relative fold change in the mRNA expression levels of cytokines in IL-1β-induced C28/I2 cells (a) and primary chondrocytes (b). Results were expressed as mean ± SD. The graphs were generated using GraphPad Prism 10.2.2.*
